# Integration of phenomics and transcriptomics data to reveal drivers of inflammatory processes in the skin

**DOI:** 10.1101/2020.07.25.221309

**Authors:** Richa Batra, Natalie Garzorz-Stark, Felix Lauffer, Manja Jargosch, Caroline Pilz, Sophie Roenneberg, Alexander Schäbitz, Alexander Böhner, Peter Seiringer, Jenny Thomas, Bentolhoda Fereydouni, Ginte Kutkaite, Michael Menden, Lam C Tsoi, Johann E Gudjonsson, Fabian Theis F, Tilo Biedermann, Carsten B Schmidt-Weber, Nikola Müller, Stefanie Eyerich, Kilian Eyerich

**Author notes:** R.B., N.G.Z., F.L. and M.J. joint first authors. N.M., S.E. and K.E. joint last authors.

## Abstract

Chronic inflammatory diseases are characterized by complex interactions between genetic predisposition and tissue-specific immune responses. This heterogeneity complicates diagnoses and the ability to exploit omics approaches to improve disease management, develop more effective therapeutics, and apply precision medicine. Using skin inflammation as a model, we developed a method that integrates deep clinical phenotyping information (phenomics) with transcriptome data of lesional and non-lesional skin (564 samples) to identify clinically-relevant gene signatures. It led us to discover so-far unexplored factors, including CCAAT Enhancer-Binding Protein Beta (*CEBPB*) in neutrophil invasion, and Pituitary Tumor-Transforming 2 (*PTTG2*) in the pathogenic epithelial response to inflammation. These factors were validated using genetically-modified human skin equivalents, migration assays, and *in situ* imaging. Thus, by meaningful integration of deep clinical phenotyping and omics data we reveal hidden drivers of clinically-relevant biological processes.

## Introduction

Ever since the emergence of high-throughput omics technologies, computational methods have played a pivotal role in biomedicine (1) (2) (3) (4) (5) (6) (7, 8). Although most of the computational algorithms can be applied ubiquitously, specific biomedical questions demand innovation. For example, identifying causative factors in chronic inflammatory processes is challenging as they are clinically heterogeneous and characterized by a complex interplay between genetic predisposition, immune responses against environmental-, self-, or microbiome-derived antigens, and tissue-specific immune response patterns of e.g. the gastrointestinal system, joints, lung, or skin. In addition and contrast to cancer, which is usually characterized by mutation-driven inexorable progression, inflammatory diseases typically show a relapsing-remitting disease course with a broad spectrum of phenotypes (9) ultimately resulting in a pronounced data variety wherein the various distinct signals are not well separated. In line with this, standardized experimental and animal models of human inflammation insufficiently reflect the comprehensive human situation (10–12), and available data sets are insufficiently characterized to dissect disease heterogeneity or identify distinct disease endotypes.

Non-communicable inflammatory skin diseases (ncISD) represent ideal prototype diseases for chronic inflammation to develop a strategy that identifies relevant molecular events within a complex network of overlaying gene expression signatures. ncISD are a group of several hundred diseases including psoriasis and atopic dermatitis (AD). They are defined by physicians using a subjective “best fit” model via investigation of a variety of attributes such as clinical phenotype, tissue and laboratory analysis (13). The transcriptome of ncISD has been widely investigated, greatly enhanced our knowledge(14) and lead to the three umbrella phenotypes which are defined according to the immune response pattern in the skin. Type 1 ncISD, characterized by cytotoxic immune cells in the skin (so-called interface dermatitis) and comprising the diseases lichen planus and cutaneous lupus(15, 16); type 2 ncISD, characterized by impaired epidermal barrier and innate immunity and comprising eczematous diseases(17, 18); and type 3 ncISD defined by Th17 immunity and the presence of neutrophil granulocytes in the skin, e.g. psoriasis and variants(19). However, the subjective nature of diagnosis, together with the heterogeneity and dynamics of ncISD, result in overlapping phenotypes and potential misdiagnoses. In addition, insufficient clinical metadata hinders discovery of relevant, potentially overlapping, disease mechanisms.

Thus, to achieve a substantial breakthrough in precision medicine, the complex pathogenesis of inflammatory diseases such as ncISD and the resulting disease heterogeneity needs to be acknowledged in research strategies. To this end, we performed deep clinical phenotyping of 287 ncISD patients using 62 attributes, including clinical picture, skin architecture, laboratory parameters, and information on the patient’s history, and generated transcriptomic profile of tissue biopsies of paired lesional and non-lesional skin. Furthermore, we developed a novel framework, called **A**ttrib**u**te **G**ene **E**xpression **R**egularization (AuGER), to integrate phenotypic/phenomic profiles with transcriptomic profiles. This approach enabled us to determine gene signatures of each clinical attribute encompassing known as well as so-far unexplored genes involved in clinically-relevant biological processes. Experimental validation of newly identified factors shed light on the drivers of neutrophil granulocyte migration into tissue and the pathogenic epithelial response to inflammation that ultimately modulates skin tissue architecture.

## Results

### Deep phenotyping explains substantially more transcriptomic tissue variance than the diagnosis alone

Given the heterogeneity in ncISD patients, we hypothesized that current clinical diagnoses only partially reflects the molecular events occurring in the skin. To test this, we collected lesional skin biopsies from 287 patients covering 13 different diagnoses of ncISD (Fig. 1A, Table S1) and performed RNA-sequencing (RNAseq). Principal component analysis (PCA) of the normalized read count of the RNAseq data showed intermixing of ncISD type 1, type 2, and type 3 diagnoses in the first two major principle components (PC1 and PC2; Fig.1A) (type 1: lichen planus, cutaneous lupus; type 2: atopic dermatitis, dishydrotic eczema, nummular eczema, hyperkeratotic-rhagadiform eczema; type 3: pustular psoriasis, guttate psoriasis, pytiriasis rubra pilaris), which indicates discrepancy between the diagnosis and the transcriptomic profile (at least in some patients). To exclude the possibility that transcriptional heterogeneity is driven by intra-individual differences, we included transcriptomic profiles of autologous non-lesional skin biopsies in the PCA (n = 564 samples in total) (Fig. S1). The status of the skin (lesional or non-lesional) explained the separation of PC1 and in total 32 % of the variance in the transcriptome, with genes related to skin inflammation driving the separation (Fig. S1A, B). Another 9 % of the variance mainly in lesional samples was explained by gender, which was confirmed by the expression of genes located on sex chromosomes (Fig. S1A, B). Thus, we performed unsupervised hierarchical clustering of lesional transcriptome excluding genes encoded on the sex chromosomes. There was still intermixing of patients with different diagnoses in the clusters (Fig. S1C), indicating that gender correction alone is insufficient to improve the concordance of transcriptome and diagnosis.

**Figure 1:**
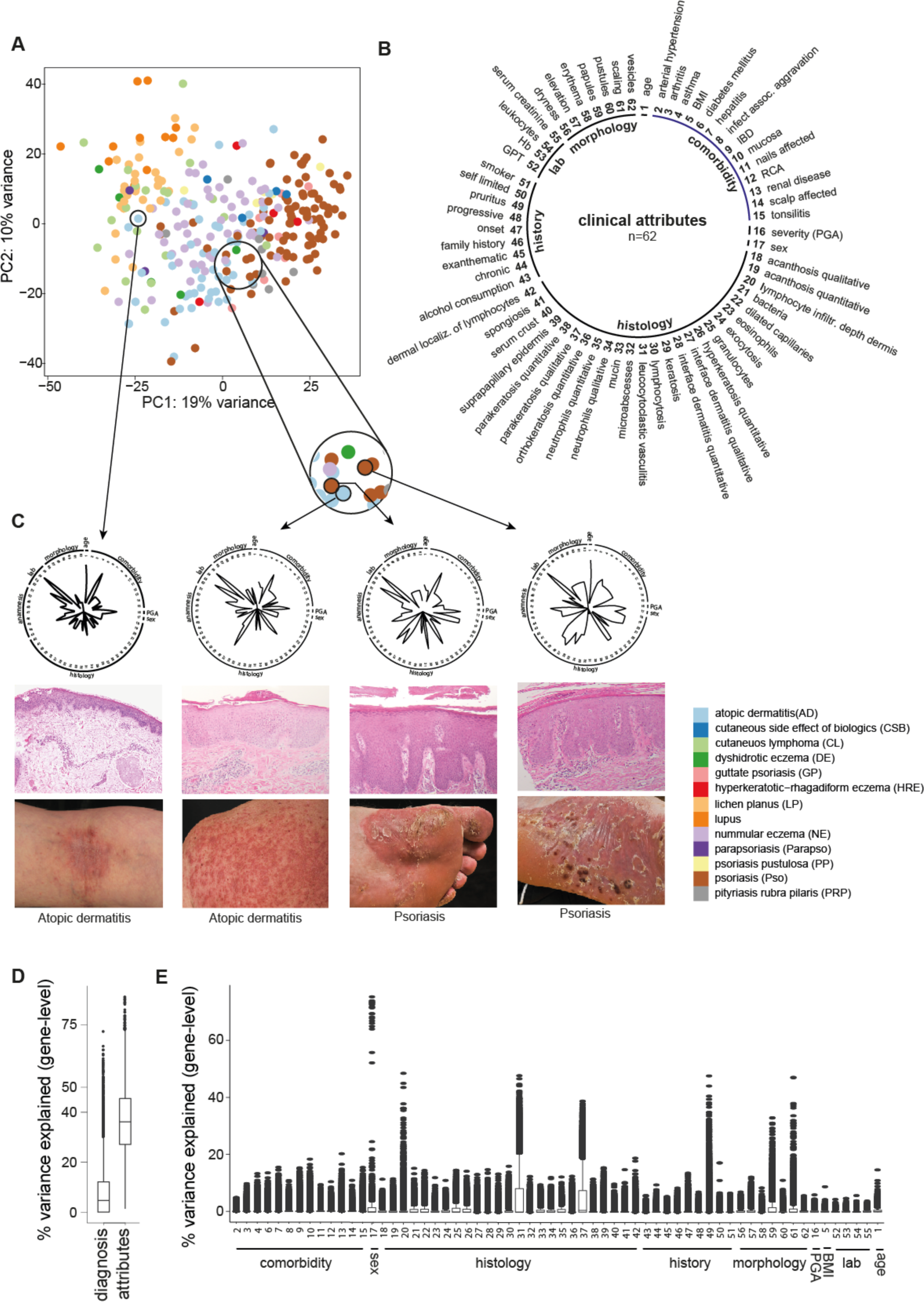
Transcriptomic heterogeneity of ncISD requires deep clinical phenotyping. A) Principal component analysis (PCA) of gene expression from 287 lesional skin biopsies, spanning 13 different non-communicable inflammatory skin diseases. Each dot represents a patient colored by its diagnosis. PCA reveals three, albeit inconcise, subclusters consisting of lichen planus/ lupus (type 1 ncISD) (light and dark orange), nummuluar eczema/ atopic eczema (type 2 ncISD) (light purple/light blue) and psoriasis (type 3 ncISD) (brown) B) Overview of clinical attributes (n=62) grouped into morphology, comorbidity, histology, patient’s history (history), and laboratory parameters (lab), used to obtain a deep clinical phenotype of each patient. C) Exemplary fingerprints of four patient’s (two atopic dermatitis and two psoriasis patients) attribute composition (radar plots), histological examination (middle) and clinical picture (right) demonstrating that two patients with a similar transcriptional profile also have a similar phenomic profile (radar plot) despite differingclinical diagnosis (patients highlighted in the circle). However, patients with the same diagnosis (e.g. atopic dermatitis as indicated) may have a heterogeneous clinical phenotype combined with transcriptional diversity. D) Transcriptional variance explained estimated using linear mixed effect model. Clinical attributes additively explain higher transcriptional variance (36.7%) compared to the clinician’s diagnosis (7.8%)E) Overview of the contribution of each individual clinical attribute to the variance in the transcriptome data set. BMI: body mass index, GPT: Glutamat-Pyruvat-Transaminase, Hb: hemoglobin, IBD: inflammatory bowel disease, RCA: rhinoconjunctivitis allergica.

To dissect transcriptional heterogeneity seen in patients, we performed deep clinical phenotyping of the 287 patients using 62 attributes from the data recorded by dermatologists during diagnosis that reflect clinical picture, histologic architecture, laboratory results, investigation of comorbidities, and personal and family history (Fig. 1B, Fig. S2). This generated 287 unique patient fingerprints (Fig. S3). We analyzed the fingerprints from four patients with the diagnosis psoriasis (n = 2) or AD (n = 2). Three of these clustered closely based on transcriptional profiles, while one AD patient was located far from the other three patients in the PCA (Fig. 1C). At this single patient level, patients with transcriptionally-similar profiles indeed represented individuals with similar phenomic profiles despite their differing diagnoses (Fig. 1C). To compare the amount of transcriptomic variance explained by attributes derived from deep clinical phenotyping with the clinicians’ diagnosis we used a linear mixed effect model. We found that clinicians’ diagnoses only explained 7.8 % of the total transcriptional variance in the skin (Figure 1D). In contrast, the 62 attributes of the deep phenotyping together explained 36.7 % of the total transcriptomic variance (Fig. 1D, E). Thus, we concluded that transcriptomic profile of a patient can be better disentangled using multiple, clinically-relevant attributes than the clinician’s diagnosis alone.

### Unique attribute-gene signatures identified by deconvolution of the transcriptional signal

Usually single attributes are not specific for a distinct disease, for instance, scaling which is considered a hallmark of psoriatic skin, can also occur in other inflammatory skin conditions. However, these overlapping attributes still contribute in shaping the lesional skin transcriptome and thereby the traditional approach of comparing diseases is limited owing to high background noise. To overcome the super-imposed definition of diseases, we evaluated each patient using several clinical attributes. With the assumption that each attribute represents a clinically-relevant biological process, we developed a computational method to identify attribute specific gene signatures. Our approach **A**ttrib**u**te **G**ene **E**xpression **R**egularization (AuGER, Fig. 2A) associates phenomic profiles with transcriptomic profiles using regularized regression. As a first step, we performed attribute reduction using pairwise correlation analysis, which revealed groups of highly correlated attributes (Fig. 2A 1-3, Fig. 2B, Table S4). As expected, attributes that are used to describe a specific ncISD disease type highly correlate with each other: specifically, the ncISD type 1 attributes, “number of dyskeratoses”, “interface dermatitis qualitative” and “quantitative”, “localization of dyskeratoses”; the ncISD type 2 attributes, “serum crusts”, and “spongiosis”; and the ncISD type 3 attributes, “neutrophils qualitative” and “quantitative”, and “parakeratosis” (Figure 2B). One representative attribute (= core attribute) per group of highly correlated attributes was selected based on clinical relevance, favouring quantitative measures over qualitative discretization, resulting in a total of 24 core attributes used for further analysis (Fig. 2C).

**Figure 2:**
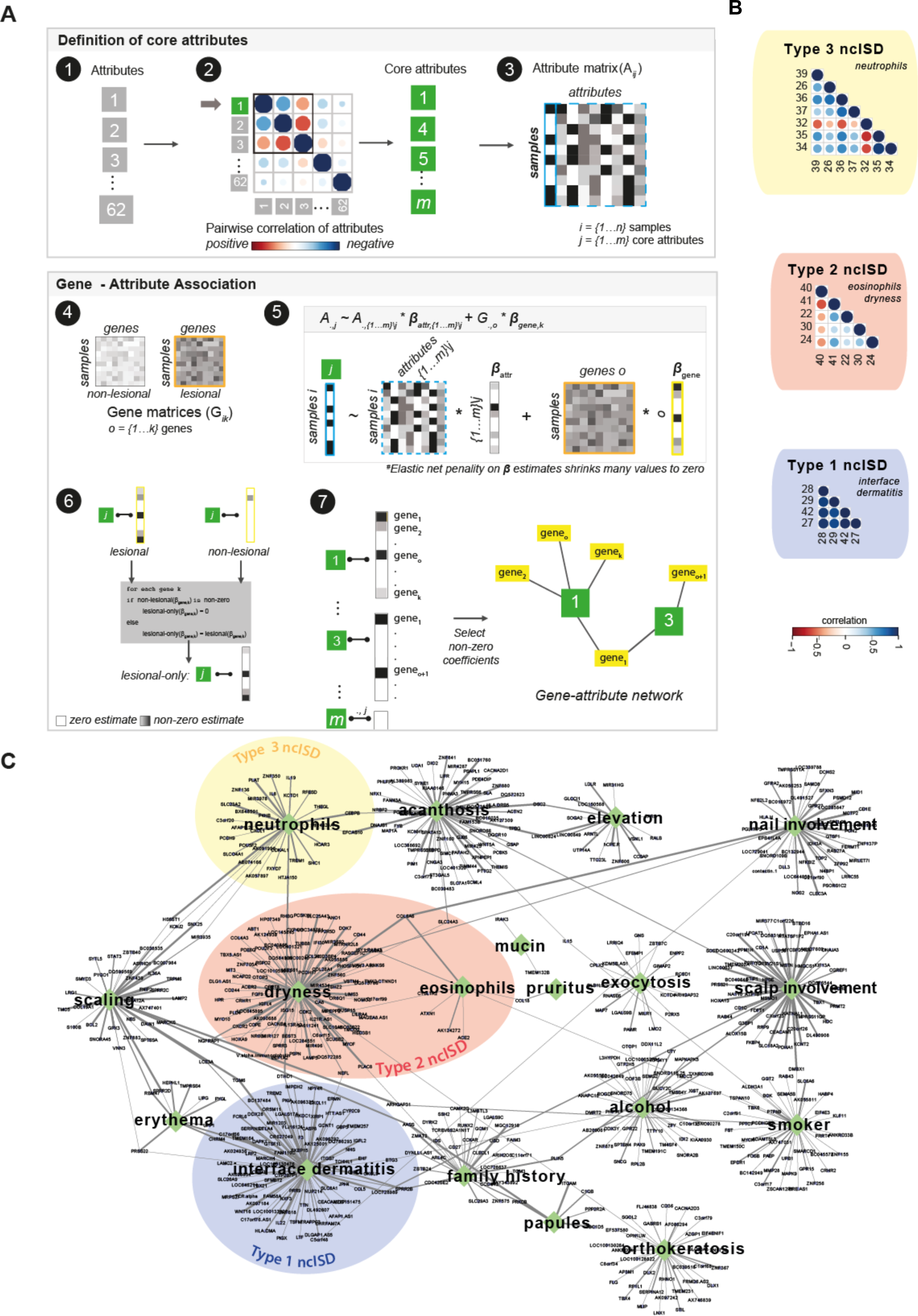
AuGER: Attribute Gene Expression Regularization. A) Schematic of the AuGER approach. 1^st^ step: Definition of core attributes. The 62 attributes of all samples *i*= {1…*n*} were usedto identify highly correlated attributes. From thehighly correlated set of attributes one representative was selected based on clinical relevance, resulting in a set of 24 core attributes *j* (*j*= {1…*m*} and in the attribute matrix A*_i,j_*. 2^nd^ step: Gene-attribute association. First, RNASeq count data was split into lesional and non-lesional sets of *k* genes (*o* = {1… *k*}, gene matrix G*_ik_*), respectively). Next, multivariate regression with elastic net penalty was used to identify the association between each core attribute (blue box) and gene expression (here illustrated for lesional skin only, orange box). The *m-1* attributes other than response were used as covariates to control for the influence on response variable (dashed blue box). Regression or beta coefficients *β_attr_* and *β_gene_* (yellow box) for each predictor variable were estimated, Then, genes with a non-zero *β_gene_* coefficient in lesional skin only were selected for each attribute and the association of genes and clinical attributes was visualized as a bipartite network. B) Exemplary highly correlated attributes belonging to type 1, type 2, or type 3 ncISD clusters with their core attribute name in italics. C) Network visualization of gene sets derived from AuGER. Attributes (green diamonds) and genes (blue ellipses) represent nodes, edge is drawn between an attribute and gene with non-zero regression coefficient, thickness of edges represent strength of an association derived from the frequency of association (see methods). Colors indicate core attributes of the three ncISD clusters

Next, we used multivariate regression with elastic net penalty to associate each core attribute with the transcriptome. For each of the 24 core attributes regression analysis was carried out, such that the respective core attribute was used as the dependent variable (output), and the genes of the complete transcriptome data as the predictor variables (input). The remaining 23 core attributes were used as covariates to control for their impact on the dependent variable. For each gene, the regression coefficient was estimated per attribute, and all genes with non-zero regression coefficients were identified (Fig. 2A 4-5). To determine the clinically-relevant associations between individual attributes and genes,, genes with non-zero coefficient in lesional and zero coefficient in non-lesional skin were selected (Fig. 2A 6). For robust results, regression analysis of both lesional and non-lesional transcriptome was conducted five times each, using five sets of imputed core attributes. Specific, unique, and shared associations were obtained between a total of 621 genes and 18 of the 24 attributes (Fig. 2A 7, Table S4, Fig. S4). For the first time, AuGER acknowledges disease heterogeneity of ncISD by dissecting diseases into multiple clinically-relevant attributes and assigning gene signatures to these attributes. These gene signatures reveal the molecular basis of complex and biologically-relevant processes, e.g. the presence of neutrophil granulocytes in the skin or pathogenic epithelial thickening as a consequence of inflammation, so-called acanthosis.

### The gene expression landscape of ncISD

To understand the interconnections of attributes and its reflection on ncISD types, we created an atlas of skin inflammation (Fig. 2C). This atlas depicts the findings of AuGER as a network, with 18 core attributes and 621 genes as nodes linked by an edge representing a non-zero regression coefficient. On the one hand, this atlas reflects core attributes that represent hallmarks of the three ncISD clusters, type 1 (lichen planus and cutaneous lupus, here identified by the core attribute “interface dermatitis”), type 2 (eczema variants which are typically associated to “dryness” and “eosinophils”), and type 3 (psoriasis variants represented by the core attribute “neutrophils”).

On the other hand, the atlas also illustrates attributes which are not typically associated with one of the three ncISD clusters, but still have an impact on the skin transcriptome. These include scalp or nail involvement, clinical presentation such as scaling, papules, or erythema; or histo-pathological presentation such as acanthosis or orthokeratosis (regular formation of the upper epidermis). These attributes may occur in several diseases and thus increase the unspecific background noise in traditional approaches used to compare different diseases. Finally, external factors influencing the lesional skin transcriptome are also captured. This is represented by numerous genes associated with the core attributes “alcohol consumption” and “smoking” (Fig. 2C).

Notably, AuGER identified only one gene associated with the attribute “pruritus”, namely the T cell recruiting chemokine *CCL18* (20). *CCL18* was shared with the core attribute “eosinophils” and thus connected to type 2 ncISD, namely eczema. This is in line with the fact that most eczema variants are highly pruritic (21). Indeed, levels of *CCL18* were highest in eczema variants (AD and nummular eczema) as compared to psoriasis (Fig. S6A). This was confirmed in an independent data set consisting of skin from healthy volunteers (n = 38) and from patients suffering from AD (n = 27) or psoriasis (n = 37). Here, expression of *CCL18* was again significantly higher in AD than in the healthy controls (log2 fold change: 4.9, p 1.4*10^-13^) or psoriasis (3.1, p 1.4*10^-7^) (Fig. S6B). In a third cohort, *CCL18* expression was higher in lichen planus lesional skin (n = 20) as compared to healthy controls (n = 38). Just like eczema, lichen planus is typically pruritic (22)(Fig. S6B).

Furthermore, all attributes except “mucin” shared at least one link with another attribute in the network, thus constructing a comprehensive gene map of skin inflammation. “Mucin” is a core attribute comprising an enhanced level of extracellular material (23). It is not associated with an inflammation cluster and is usually the result of inflammation of deeper skin layers rather than of the epidermis. Connections of the individual gene signatures of each attribute were also identified at the level of pathway analysis. Some pathways were associated with only one attribute, e.g. the expected cytotoxicity-mediating pathway “IFN-gamma signaling” associated with the ncISD type 1 core attribute “interface dermatitis” (15). Numerous pathways were associated with at least five different attributes. For example, pathways of cell cycle and proliferation, such as “EGF signaling”, “Ras signaling”, “Oncostatin M signaling”, and the stem cell factor “Wnt signaling”, connected the clinical attributes “elevation” (palpable skin lesions) and “papules”, both of which may be related to exaggerated proliferation of the epidermis (Fig. S5).

### Novel insights into neutrophil granulocyte biology in tissue

The comprehensive atlas of the transcriptome landscape of ncISD delivers a plethora of so-far undiscovered connections of clinically-relevant biological processes with gene expression. To understand the pathogenicity of these genes, we explored the genes associated with the attribute “neutrophils”, which was defined as the presence of neutrophil granulocytes in lesional skin as observed by dermatopathology. AuGER identified an association of 35 genes with this attribute (Fig. 3A). While some of these genes such as *CXCL1* or *CXCL8* are well-known drivers of neutrophil migration and/or biology (24), others were previously unknown in this context. As expected, the ncISD type 3 core attribute “neutrophils” was most prominent in psoriasis patients, however, intra- and inter-disease variability was high (Fig. 3B, C). Of the 35 genes majority are associated with inflammation and general processes such as protein binding (Fig. 3D). For biological validation of the attribute-gene association, we focused on five genes *CEBPB*, *HCAR3*, *SHC1*, *HS6ST1* and *SLC23A2* as they were centrally located in the network (Fig. 2C) and are novel in the context of skin diseases. First, we analyzed the effect of stimulation with pro-inflammatory cytokines *IL-17A*, *TNF-α* or *LPS* on gene expression in two potential target cells, primary human keratinocytes and neutrophil-like HL-60 cells. Stimulation induced gene expression only of *CEBPB* consistently in both cell types as compared to unstimulated controls (Fig. 4A).

**Figure 3:**
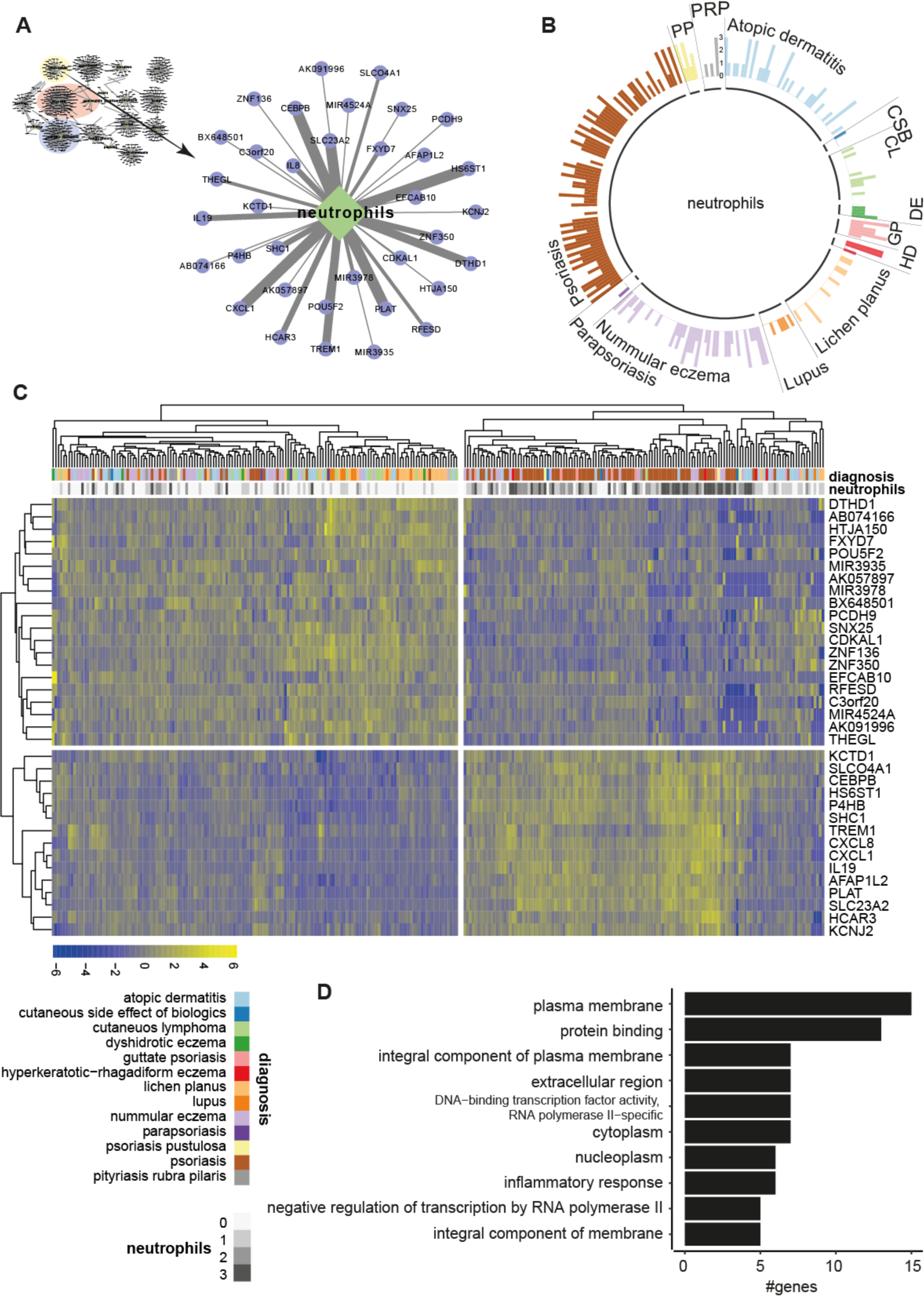
The attribute ‘neutrophils’. A) A focus on the clinical attribute ‘neutrophils’ and its associated genes. Line thickness indicates the strength of gene-attribute associationB) Distribution of the attribute ‘neutrophils’ across the 13 non-communicable inflammatory skin diseases. Each bar represents one patient, and the height of the bar indicates the number of neutrophils present in skin histology with 0 = no, 1 = low, 2 = moderate, 3 = high numbers of lesional neutrophils (see Fig. S2). C) Hierarchical clustering of neutrophil-associated genes annotated with diagnosis (colors) and presence of neutrophils (grayscale). D) Gene ontology terms associated with the neutrophil gene set. CL: cutaneous lymphoma; CSB: cutaneous side effect of biologics; DE: dishydrotic eczema; GP: guttate psoriasis; Parapso: parapsoriasis; PP: psoriasis pustulosa; PRP: pityriasis rubra pilaris.

**Figure 4:**
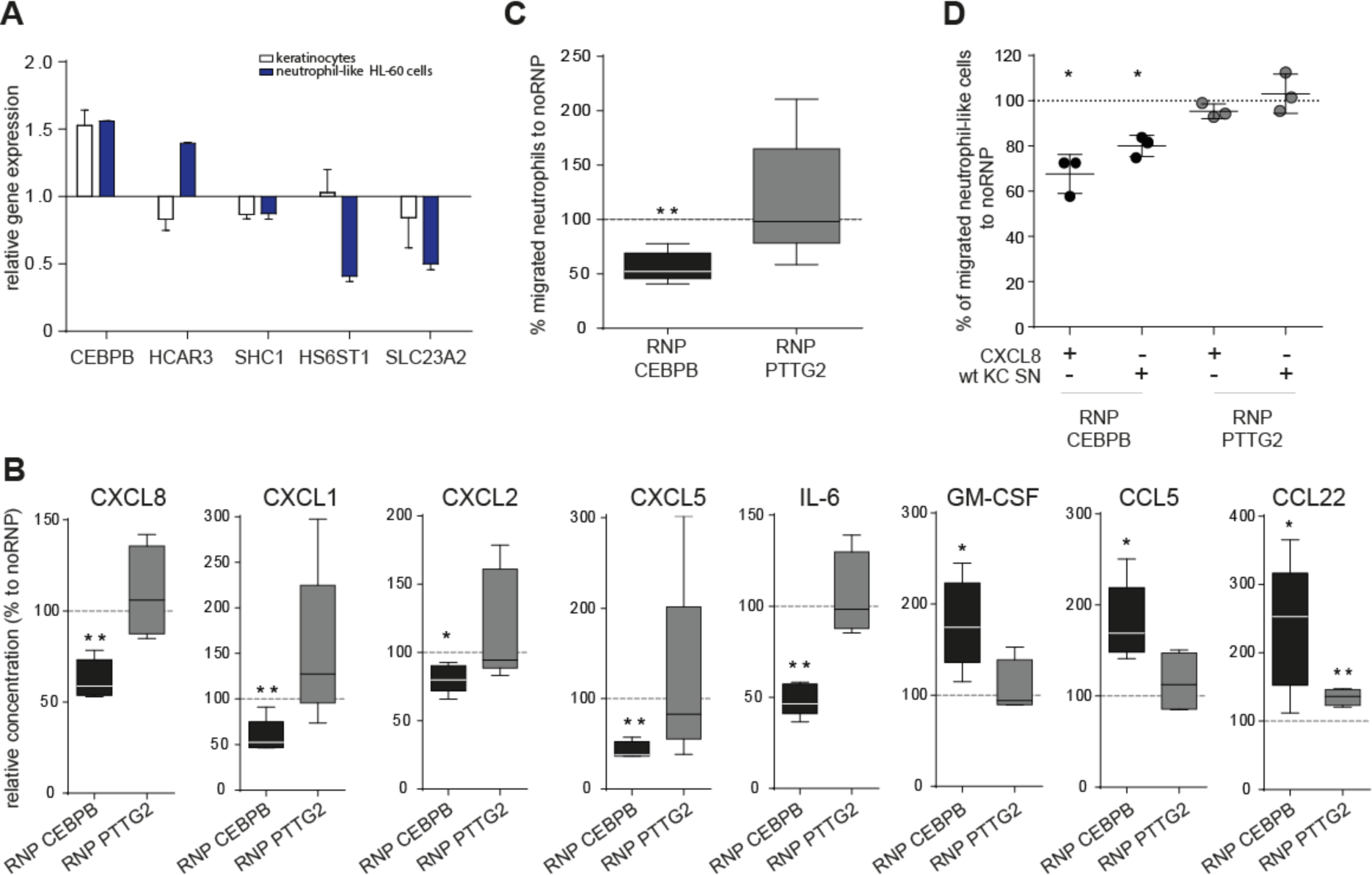
CEBPB regulates neutrophil granulocyte biology. A) Five genes with a strong association to the attribute ‘neutrophils’ and a yet unknown function in this field were selected and their relative gene expression was evaluated in primary human keratinocytes (stimulated with IL-17A and TNF-α) and neutrophil-like cells (differentiated HL60 cells) (stimulated with LPS) compared to non-stimulated controls by real time PCR. B) *CEBPB* and *PTTG2* were knocked out in primary human keratinocytes using the RNP-CRISPR technology. Knock-out (RNP) and wild-type (no-RNP) keratinocytes were cultured and stimulated with recombinant *IL-17A* for 72 hours to induce neutrophil-relevant genes. The supernatant was analyzed for cytokine/chemokine content by multiplex technology and relative changes are given in the graphs as percentage compared to no-RNP controls. C) Frequency of neutrophils migrated towards the supernatant of keratinocytes stimulated with recombinant *IL-17A* after *CEBPB* or *PTTG2* knock-out compared to *IL-17A*-stimulated wild type keratinocytes. D) Knockout of *CEBPB* or *PTTG2 in* neutrophil-like cells (differentiated HL60 cells) and consecutive migration towards *CXCL8* or supernatant of wild type keratinocytes (wt KC SN) stimulated with recombinant *IL-17A*. KC: keratinocyte; RNP: ribonucleoprotein; SN: supernatant.

C/EBP factors, to which *CEBPB* belongs, have been found to regulate cytokine expression in human neutrophils (25). To gain more insight into the function of *CEBPB* in neutrophil biology in particular into the interaction of neutrophils and keratinocytes, we specifically knocked-out *CEBPB* and the unrelated control gene *PTTG2* in keratinocytes using CRISPR/Cas9 technology (controls on knock-out efficiency are given in Fig. S7). Only *CEBPB* knock outs (not *PTTG2*) resulted in significant reduction of central chemokines for neutrophil migration and activation (*CXCL8, CXCL1, CXCL2, CXCL5, IL-6*) in the supernatant of keratinocytes stimulated with *IL-17A* (Fig. 4B). Moreover, *CEBPB* knock-out cells produced significantly higher amounts of the growth factor *GM-CSF* and the chemokines *CCL5* and *CCL22*, indicating that loss of *CEBPB* does not hamper the ability of keratinocytes to promote bone marrow neutrophil maturation, or migration of eosinophils, macrophages or dendritic cells (Fig. 4B). To validate *CEBPB* mediated regulation of neutrophil migration to the skin, we quantified migration of primary human neutrophils towards the supernatant of *CEBPB* or *PTTG2* knock-out keratinocytes, compared to the supernatant of wild type controls. Here, knock-out of *CEBPB* significantly reduced the capacity of keratinocytes to regulate neutrophil migration (Fig. 4C). Furthermore, knock-out of *CEBPB* directly in the neutrophil-like HL-60 cells resulted in diminished migration towards both *CXCL8* and keratinocyte supernatant (Fig. 4D). Taken together, we identify *CEBPB* is a pivotal regulator of neutrophil biology in the skin, acting at both the keratinocyte and the neutrophil level.

### Pathogenic epidermal hyperplasia is driven by previously unknown genes

Pathogenic hyperplasia of the epidermis caused by excessive proliferation and disturbed differentiation of keratinocytes, so-called acanthosis, is a process frequently observed in different kinds of skin inflammation. Therefore, acanthosis is a representative example of AuGER’s power to select and identify so-far undiscovered drivers of this highly prevalent clinical phenotype. 69 genes were associated with acanthosis (Fig. 2C, Fig. 5A). Acanthosis is neither specific for a ncISD type, nor is it observed in all patients suffering from a ncISD. In line with this, in our cohort, acanthosis as an attribute was variably distributed across the 13 types of ncISD (Fig. 5B) and unsupervised hierarchical clustering of genes associated with acanthosis in all 287 patients did not correlate with distinct diseases (Fig. 5C). The acanthosis gene signature is associated with cell-cell adhesion and activity (Fig. 5D).

**Figure 5:**
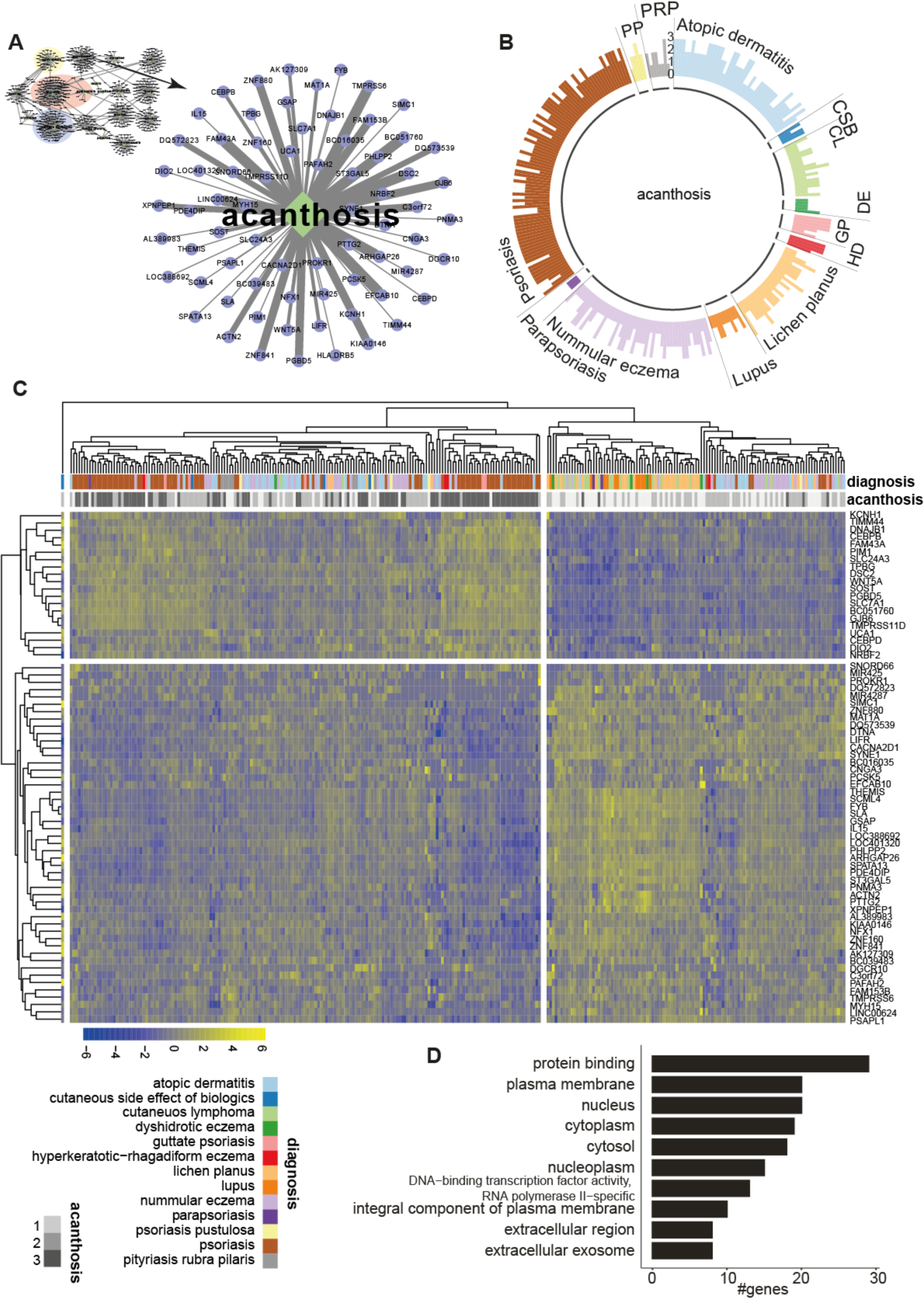
The attribute ‘acanthosis’. A) A focus on the clinical attribute ‘acanthosis’ and its associated genes. Line thickness indicates the strength of the gene-attribute association. B) Distribution of the attribute ‘acanthosis’ across the 13 non-communicable inflammatory skin diseases. Each bar represents one patient, and the height of the bar indicates the level of acanthosis measured by skin histology, with 0 = no, 1 = low-, 2 = moderate, 3 = strong acanthosis (see also Fig. S2). C) Hierarchical clustering of acanthosis-associated genes annotated with diagnosis (colors) and strength of acanthosis (grayscale). D) Gene ontology terms associated with the acanthosis gene set. CL: cutaneous lymphoma; CSB: cutaneous side effect of biologics; DE: dishydrotic eczema; GP: guttate psoriasis; Parapso: parapsoriasis; PP: psoriasis pustulosa; PRP: pityriasis rubra pilaris.

For biological validation of the attribute-gene association, we selected *PTTG2*, *SYNE1*, and *CEBPB*. Keratinocytes stimulated with the pro-inflammatory cytokines *IL-17A* and *TNF-α* showed an upregulation of *CEBPB* and a downregulation of *SYNE1* gene expression, while *PTTG2* expression remained unaltered (Fig. 6A). Cellular morphology or the architecture of three-dimensional human skin equivalents remained unaltered in knock-out of *PTTG2*, *SYNE1*, *CEBPB* in primary human keratinocytes (Fig. 6B, see effectiveness in Fig. S7). However, it is difficult to judge on specific effects in unstimulated models as they only consist of a few cell layers. The cytokine IL-22 is a known inducer of keratinocyte growth and induces acanthosis (26). Next, we investigated the effects of *CEBPB*, *SYNE1* and *PTTG2* knock-out on epidermal thickness as a proxy for acanthosis in response to *IL-22*. In line with the differences in gene regulation, knock-out of both *PTTG2* and *CEBPB* significantly inhibited the induction of acanthosis by *IL-22* compared to the control (noRNP) model, with *CEBPB* preventing it completely. In contrast, knock-out of *SYNE1* markedly increased the thickness of the three-dimensional skin model (Fig. 6B, 6C; Fig. S8). Notably, proliferation of keratinocytes was also reduced by knock-out of *CEBPB* or *PTTG2* but induced by *SYNE1* (Fig. 6C, 6D, 6E). In addition, knock-out of *CEBPB* resulted in altered epithelial differentiation after *IL-22* stimulation, namely in clearly diminished expression of the early differentiation marker keratin 10, and keratin 16, which is typically associated with proliferative keratinocytes (27) (Fig. 6E). Hence, *CEBPB* and *PTTG2* are essential for the development of acanthosis, while *SYNE1* is a negative regulator of epidermal thickness.

**Figure 6:**
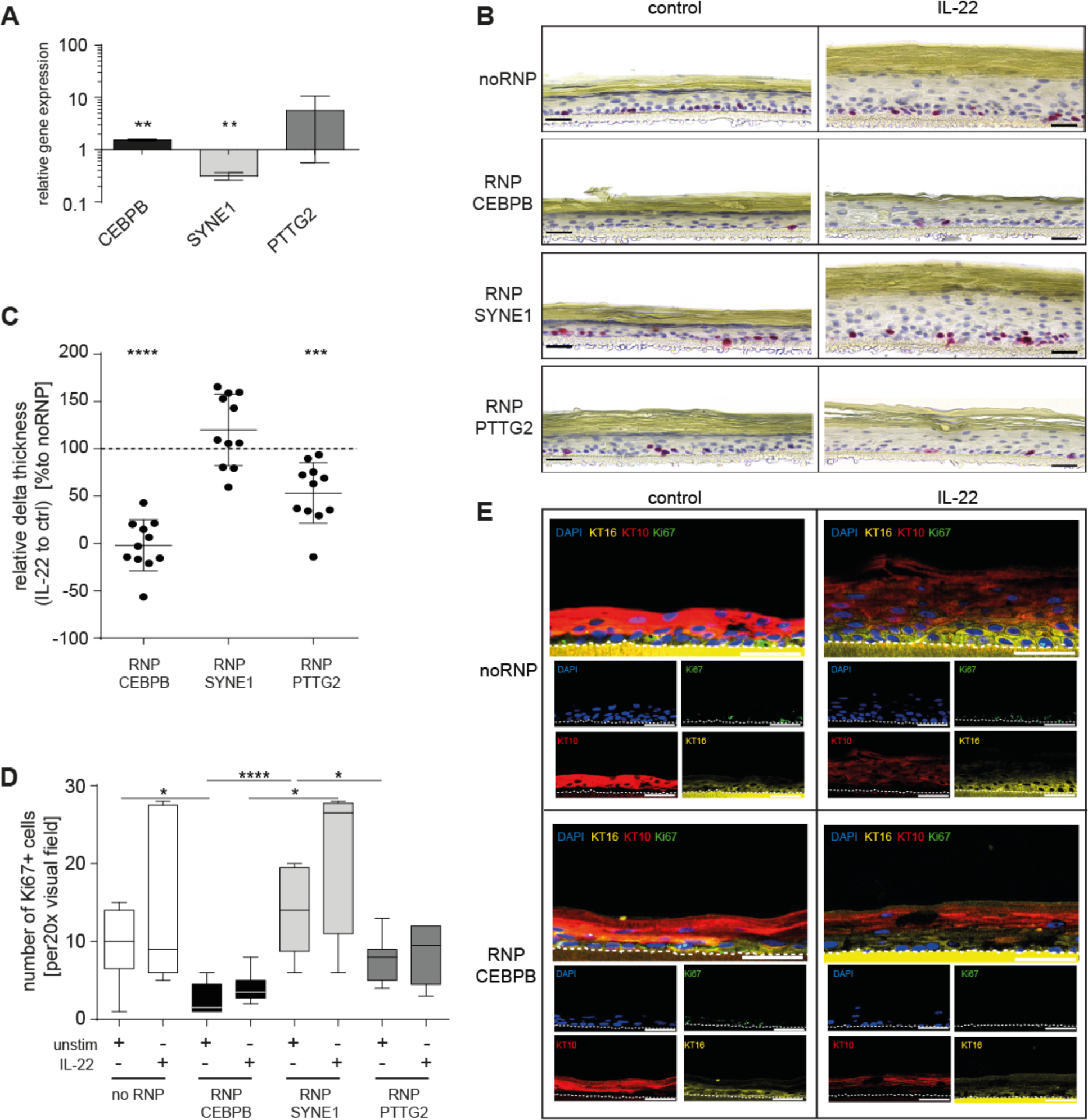
Validation of acanthosis-related genes. A) Genes with a strong and yet unknown association to the attribute acanthosis were chosen and their expression was evaluated in primary human keratinocytes after *IL-17A* and *TNF-α* stimulation by real time PCR. B) The acanthosis-associated genes *CEBPB, SYNE1 and PTTG2* were knocked out in primary human keratinocytes using CRISPR-Cas9. 3D skin models of knock-out (RNP) and wild-type (no-RNP) were stimulated with recombinant *IL-22* to induce acanthosis or left untreated (control) and stained for the proliferation marker Ki67. Representative staining are shown. Bars represent 40µm. C) Relative delta thickness of 3D skin models with RNP-CRISPR knock-out for *CEBPB*, SYNE1 or *PTTG2* (n = 11). Here, thickness of each *IL-22* stimulated model was normalized to its untreated control followed by calculation of percental changes of RNP vs no-RNP treated models. D) Quantification of Ki67 positive cells in 3D skin models shown in B). E) Representative Keratin 10 (KT10) (red), Keratin 16 (KT16) (yellow), and Ki67 (green) immunofluorescence staining of 3D skin models with a RNP-CRISPR knock out of *CEBPB* with or without (control) IL-22 stimulation. Nuclei were stained with DAPI (blue). The dotted white line indicates the border between basal keratinocytes and the culture membrane. Results on *SYNE1* and *PTTG2* are given in Fig. S8. Bars represent 50 µm. KT: keratin

## Discussion

Here, we show that so-far unexplored drivers of key pathogenic events in human inflammatory diseases can be identified from transcriptomics when disease heterogeneity is acknowledged by taking deep clinical phenotyping into account. We investigated non-communicable inflammatory skin diseases (ncISD) as a prototype example of heterogeneous human inflammatory disorders.

We observed that clinician assigned diagnosis explained very little transcriptomic variation. In contrast, metadata with a higher granularity of the observed clinical heterogeneity explains a substantially higher proportion of the transcriptomic variability associated with inflammation. Furthermore, we demonstrate that common hallmarks of the interplay between skin epithelia and immune cells are shared between different patient cohorts independent of the disease background. We and others recently followed this approach for ncISD that share the histological hallmark “interface dermatitis”, which represents cytotoxic activity of lymphocytes at the basal membrane of the epidermis and is frequently observed in type 1 ncISD such as lichen planus and lupus (15, 16). Here, identification of the molecular mechanism, type 1 immune mediated epithelial apoptosis and necroptosis, led to potential novel therapeutic strategies (28). Our current analysis takes into account other dependent clinical attributes and gives an even clearer picture of the genes involved in interface dermatitis.

Even in settings that are heavily researched and where numerous drivers are known, such as neutrophil biology, assigning more granular metadata of clinical phenotyping identifies so-far unexplored, but important mechanistic factors. In particular, our newly developed algorithm AuGER identified CEBPB as a master regulator of neutrophil biology in the skin. C/EBP proteins are known to be involved in cytokine production such as *CXCL8* and *CCL3* by neutrophils (25, 29). Here, we show that *CEBPB* is also expressed in keratinocytes, and knock-out of *CEBPB* in primary human keratinocytes results in reduced chemokine secretion and reduced neutrophil migration. This newly identified dual mechanism of action to inhibit neutrophil inflammation in tissue renders *CEBPB* a potentially powerful therapeutic target for neutrophilic inflammatory diseases of the skin, such as psoriasis or pyoderma gangrenosum but also for diseases beyond the skin as for example reactive arthritis or neutrophil-rich chronic obstructive lung disease (COPD).

Our approach not only reveals the most significant genes contributing to an attribute, but *vice versa* also identifies the mechanistic significance of single genes for corresponding attributes representing complex mechanistic processes. This was proven for the less restricted attribute “acanthosis”. Acanthosis corresponds to reactive thickening of the epidermis in inflammatory conditions and it can be observed in multiple ncISD but is not part of a disease definition (30). *PTTG2*, a hub gene for acanthosis in our analysis, has not been proposed in the context of acanthosis before. *PTTG2* is involved in cell viability and migration by regulating the expression of vimentin and E-cadherin and overexpression of *PTTG2* was described in psoriatic epidermis (31, 32). We now show that *PTTG2* is directly involved in acanthosis, as inhibiting this single factor significantly impaired acanthosis formation *in vitro*. *SYNE1*, another gene significantly correlated to acanthosis from our network, encodes an important scaffold protein. In genetically modified mice, loss of *SYNE1* leads to thickening of the epidermis (33), an observation we validated in the human system by showing increased thickness of three-dimensional skin equivalents comprised of *SYNE1* knocked-out keratinocytes. Interestingly, *CEBPB*, associated with the attribute “neutrophil” was also associated with “acanthosis” and the third candidate gene we validated for its contribution to acanthosis *in vitro*. In fact, human skin equivalents of keratinocytes lacking *CEBPB* were vital and did not show any morphological alterations, but they did not respond to *IL-22*, an inflammatory stimuli that normally induces acanthosis (26). Hence, given a tolerable safety profile, *CEBPB* could be a target to address diseases in which neutrophils and acanthosis are concomitantly present, in particular in psoriasis, hidradenitis suppurativa, or severe acne.

Our *proof-of-concept* study has limitations. First, phenotyping was partially based on subjective information, such as the patient’s narrative, clinical examination and histological evaluation. Second, though we performed RNA-Sequencing of 287 patient’s tissue samples, the sample size is still limited, and underestimation of real biological effects cannot be ruled out. Higher sample size and/or additional datasets are required to strengthen our approach, AuGER, and its potential to extract markers predictive of medically relevant outcomes, such as response to therapy. AuGER is optimized to identify genes so-far hidden in the background noise, by correcting each attribute signature for other attributes, thus, gene signatures may seem incomprehensive. Despite the limitations, this study clearly shows that associating granular clinical phenotypes to omics datasets critically determines the quality of outcome of the analysis. Phenotyping methods such as image-based machine-learning as well as adding several molecular information sources such as serum proteome or microbiome (34)will further improve our novel approach in the future.

Taken together, exemplifying ncISD we successfully combined deep clinical phenotyping with transcriptome and established that this novel approach has the potential to define pathogenic events in complex processes such as chronic human inflammation and thus build the basis for precision medicine in the field.

## Methods

### Data, Code and Material Availability

Available upon request. This study did not generate new unique reagents.

### Study cohort and deep clinical phenotyping

Patients with ncISD (n = 287, 13 distinct diagnoses, male n=163, female n=124, age male 54,19±16,69 years, age female 57,92±18,82) from the *Biobank Biederstein* were enrolled into the study. All patients were investigated by dermatology specialists. A washout phase of at least two weeks for topical treatments and five half-life times for systemic treatments was mandatory for study inclusion. The study followed the Declaration of Helsinki and was approved by the local ethics committee (Klinikum Rechts der Isar, 44/16 S). The *Biobank Biederstein* follows data protection rules and is approved by the local ethics committee (Klinikum Rechts der Isar, 5590/12). After written informed consent was obtained from all patients, each patient was deep-phenotyped regarding clinical presentation (distribution and phenotype of lesions), personal and family history (questionnaires), and laboratory parameters (smear test cultures and serum analysis). Furthermore, a non-lesional and a lesional skin biopsy (6 mm each) were obtained under local anesthesia from all patients except for ten patients that did not have non-lesional skin as all skin was affected by inflammation. Biopsy specimens were divided and one part was used for hematoxylin-eosin staining from paraffin-embedded skin. These sections were evaluated by a dermato-pathologist in a blinded manner. A list of all parameters evaluated is given in Fig. S2. Effects of sex were evaluated, reported in the results section and the corresponding figures Fig. 1 and Fig. S1.

### Primary human keratinocytes

Primary human epidermal keratinocytes (male n=4, female n=1) were obtained by suction blister as reported previously (*10*) and cultured in keratinocyte medium (DermaLife Basal Medium supplemented with DermaLife K LifeFactor Kit (Lifeline Cell Technology, LL-0007)) at 37 °C, 5 % CO_2_. Second-to third-passage primary human epidermal keratinocytes were used for all experiments.

### Primary neutrophils and neutrophil-like cells (HL-60 cells)

Primary human neutrophils were isolated from peripheral blood of healthy donors (male n=1, female n=2). HL-60 cells represent a promyelocytic cell line with phagocytic and chemotactic functions that has been obtained from a 36-year old Caucasian female with acute promyelocytic leukemia.

### RNAseq library preparation, sequencing, mapping and quantification

RNA from skin biopsies was isolated using the QIAzol Lysis Reagent (Qiagen) and miRNeasy Mini Kit (Qiagen) according to manufacturer’s protocol. RNASeq libraries were generated using the TruSeq Stranded Total RNA Kit (Illumina) according to manufacturer’s high sample protocol. Finally, samples were sequenced on an Illumina HiSeq4000 as paired-end with a read length of 2x 150 bp and an average output of 40 Mio reads per sample and end.

Sequence alignment was performed using STAR aligner with human genome reference hg38. RNAseq count data was normalized and then transformed using variance stabilizing transformation from the bioconductor package DESeq2. In total, 564 samples (287 lesional, 277 non-lesional) samples were used for further analysis.

### Principal component analysis (PCA)

Principal component analysis was performed using the top 1000 most varying genes across samples. Variance was computed per gene across the samples and 1000 genes with highest variance was used in the principal component analysis. PCA was performed twice, once for all 564 samples and once for 287 lesional samples only.

### Variance estimation

To estimate the variance in lesional gene expression explained by differences in diagnosis, gender, age and other clinical attributes, the RNAseq count data was first transformed using voom transformation from R package Limma and then modelled with a linear mixed effect model using bioconductor package variancePartition.

### Attribute-Gene Expression Regularization (AuGER)

**Step1:** Define core attributes - A total of 62 clinical attributes (Fig. S2) from 287 patients/samples were used for this analysis. Missing values (<= 20 %) were imputed using the R package mice. To identify highly correlated clinical attributes, pairwise comparison between attributes was performed using the hetcor function from the R package polycor. This function computes pearson correlation between numeric variables, polyserial correlation between numeric and ordinal variables, and polychoric correlation between ordinal variables. Of the highly correlated attributes, a representative was selected based on clinical relevance for the follow up analysis, leading to a set of 24 core attributes. The core attributes were reimputed generating 5 sets. **Step 2:** Transform the gene expression data – RNASeq count data was split into lesional and non-lesional sets. Both the sets were transformed using variance stabilizing transformation. 10 samples did not have a matching non-lesional sample. These missing samples were imputed with the median value of each gene, of non-lesional samples. **Step 3**: Regularization - Multivariate regression with elastic net penalty was used to identify the association between clinical attributes and gene expression. The regression was performed using R package glmnet, alpha was set to 0.5, lambda was computed using five-fold cross validation, family of regression was selected based on the response variable (binomial for binary, gaussian for numeric and multinomial for categorial variables). Each clinical attribute was used as response variable and gene expression data was used as predictor variables. The attributes other than response were used as covariates to control for the influence on response variable. Clinical attributes age and gender were accounted for in each regression. This step was repeated 5 x 2 times, for 5 imputed clinical attribute sets and the 2 gene expression sets of lesional and non-lesional skin. **Step 4:** Selection of genes –A regression/beta coefficient for each predictor variable was estimated by the regression analysis. All genes with a non-zero coefficient were selected for each attribute from all ten regression sets, however, genes unique to lesional skin were selected for further analysis i.e genes with non-zero coefficient in non-lesional skin were discarded. Next, genes were ranked according to the frequency of their detection in the five sets of lesional skin. We refer to these ranks as strength of association between an attribute and the gene. **Step 5:** Network – The association of genes and clinical attributes was visualized as bipartite network. Genes and clinical attributes were represented as nodes and edges/links between them indicate associations. The edge/link width indicates the rank of the genes. The network was visualized using cytoscape. Nodes were manually arranged to aid the visualization.

### Culture of primary human keratinocytes, primary neutrophils and neutrophil-like cells (HL-60 cells)

Primary human epidermal keratinocytes were cultured in keratinocyte medium (DermaLife Basal Medium supplemented with DermaLife K LifeFactor Kit (Lifeline Cell Technology)) at 37 °C, 5 % CO_2_. Second-to third-passage primary human epidermal keratinocytes were used for all experiments. Before stimulation, cells were starved for 5 hours in DermaLife Basal Medium followed by stimulation with human recombinant IL-17A (R&D systems, 50 ng/ml) and TNF-α (R&D systems, 10 ng/ml) for 16 hours (mRNA analysis) or 72 hours (production of supernatant) in keratinocyte medium without hydrocortisone.

Primary human neutrophils were isolated from peripheral blood of healthy donors using LymphoPrep™ (Progen) and 2 % Dextran solution (Sigma Aldrich) and a final erythrocyte lysis. Neutrophils were cultured in RPMI 1640 medium supplemented with 1 % human serum, 0.1 mM NEAA, 2 mM L-Glutamine, 1 mM sodium pyruvate and 100 U/ml penicillin/streptomycin at 37 °C, 5 % CO_2_.

HL-60 cells were cultured in HL-60 basal medium (RPMI1640 + HEPES & GlutaMAX (GIBCO) supplemented with 9 % FCS and 100 U/ml penicillin/streptomycin) at 37 °C, 5 % CO_2_. To obtain neutrophil-like cells, HL-60 cells were differentiated for 6 days in HL-60 basal medium supplemented with 1.3 % DMSO (PanReac & AppliChem ITW Reagents) as published before (35). For gene expression analysis HL-60 cells were stimulated with 100 ng/ml LPS (InvivoGen) for 6 hours.

### Neutrophil migration assay

Migration assay was performed using ChemoTx® Disposable Chemotaxis System (neuroprobe). 3 x 10^4^ neutrophils were added to the top of a 5 µm pore polycarbonate membrane and migrated to keratinocyte supernatant for two hours. Migrated cells were analysed with an LSRFortessa flow cytometer (BD Biosciences). Migration was performed in triplicates.

### CRISPR-Cas9 knock-out

**Preparation of RNP complexes:** Predesigned crRNA, tracrRNA, tracrRNA-ATT and S.p. HiFi Cas9 Nuclease was ordered from Integrated DNA Technologies in Alt-R^®^ format. CrRNA and tracrRNA was reconstituted to 200 µM with IDTE Buffer. For generation of crRNA-tracrRNA duplex Oligos were mixed at equimolar concentrations and annealed by heating at 95 °C for 5 min following a slow cool-down to room temperature. For each target, a mixture of two crRNA sequences was applied. Finally, for RNP formation 180 pmol crRNA-tracrRNA duplex was mixed with 60 pmol Cas9 protein in Nucleofection Buffer P3 (Lonza) and incubated for at least 10 min at room temperature. crRNA sequences can be found in Table S2. **Transfection of RNP complexes:** CRISPR knockout was done by application of RNP complexes with the 4D Nucleofector™ device (4D-Nucleofector™ Core Unit; 4D-Nucleofector™ X Unit, both Lonza) using the P3 Primary Cell 4D-Nucleofector™ X Kit S (Lonza) for keratinocytes or the SF Cell Line 4D-Nucleofector™ X Kit S (Lonza) for HL-60 cells. For transfection, 1.6 x 10^5^ keratinocytes or 1 x 10^6^ HL-60 cells were resuspended in 16 µl Nucleofection Buffer P3 (for keratinocytes) or SF (for HL-60 cells) and mixed with 4 µl RNP complex. The transfection mixture was transferred to Nucleofection™ cuvette strips and electroporated with the impulse DS-138 (for keratinocytes) or EO-115 (for HL-60 cells) in the 4D Nucleofector™ X unit. After nucleofection, cells were rested at room temperature for 10 min. Afterwards, 80 µl prewarmed cell culture media was added and cells were transferred to cell culture plates of choice depending on application.

### 3D keratinocyte skin model

3D keratinocyte skin models were cultured in polycarbonate inserts (Millipore) placed in a tissue-culture treated 6-Well plate. Inserts was pre-coated with 1 % collagen type I solution (SIGMA) in PBS for 45 min. 0.3 x 10^6^ primary human keratinocytes were seeded directly after transfection in 500 µl keratinocyte medium supplemented with 1.5 mM CaCl_2_ (SIGMA) into the insert. 2.5 ml keratinocyte medium supplemented with 1.5 mM CaCl_2_ was added in the surrounding well. The models were cultured at 37 °C and 5 % CO_2_. Two days after seeding, airlift was done by aspirating the medium in the insert and the medium in the surrounding 6-well was replaced with 1.8 ml keratinocyte medium supplemented with 1.5 mM CaCl_2_ and 50 µg/ml Vitamin C (SIGMA). Every second day the medium was exchanged with 2 ml keratinocyte medium supplemented with 1.5 mM CaCl_2_ and 50 µg/ml Vitamin C in the surrounding 6-well. Nine days after airlift, 3D keratinocyte skin models were stimulated with 50 ng/ml IL-22 (R&D) in 2 ml keratinocyte medium supplemented with 1.5 mM CaCl_2_ and 50 µg/ml Vitamin C and without hydrocortisone for three days. For histological analysis, inserts were fixed with 4 % Formaldehyde for 24 h at 4 °C and embedded in paraffin. The epidermal thickness of 3D skin models was measured from the top of the corneal layer to the bottom of the basal keratinocyte layer with ImageJ (https://imagej.nih.gov/ij/download.html) in four sections with a distance of 100 µm each per sample and the mean was calculated. Next, delta thickness (IL-22 minus unstimulated thickness) was calculated for each knockout group to visualize the acanthosis effect. Finally, this delta thickness was illustrated in relation to the delta thickness from the untransfected noRNP control sample.

### Immunohistochemistry

5 µm sections of 3Dskin model samples were air-dried overnight at 37 °C, dewaxed and rehydrated. Staining were performed by an automated BOND system (Leica) according to the manufacturer’s instructions: After antigen retrieval in a pH9 EDTA buffer based epitope retrieval solution (Leica), sections were incubated with monoclonal antibodies against Ki67 (rabbit anti-Ki67, Zytomed), and a secondary polymeric alkaline phosphatase (AP)-linked anti-rabbit antibody was applied. The complex was visualized by the substrate chromogen Fast Red. Slides were counterstained with haematoxylin. As a negative control, primary antibodies were omitted. Positive cells were counted in four to nine visual fields per condition.

### Immunofluorescence

Paraffin mounted sections were air-dried overnight at 37 °C and dewaxed at 65 °C for 25 minutes and rehydrated by consecutive washes in X-TRA-Solv® (Medite), isopropanol, ethanol (96 % vol/vol and 70 % vol/vol, respectively) and distilled H_2_O. Antigen retrieval was performed in a pressure cooker with boiling citrate buffer (pH6). Hydrogen peroxide (Sigma Aldrich, Germany) blocking was performed in 3 % solution for 15 minutes. Sections were further blocked with 10 % normal goat serum and 10 % normal donkey serum diluted in antibody diluent (Abcam) for one hour and incubated with primary antibody mix (anti-ki67, anti-KT10 antibody & anti-KT16 antibody, all Abcam) for one hour at room temperature and then overnight. For negative controls, antibody diluent only was applied. After overnight incubation, secondary antibodies (AF488 donkey anti-rat, AF647 goat-anti rabbit antibody & AF557 donkey anti-mouse antibody, all Thermo Fisher Scientific) were applied in the dark for one hour at room temperature. Sections were treated with Vector® TrueVIEW® autofluorecence quenching kit (VECTOR Laboratories) and mounted with the included VECTASHIELD® Vibrance™ plus DAPI (VECTOR Laboratories). Stainings were imaged in the blue (DAPI), green (KI67), red (KT10) and far red (KT16) channel of a Leica TCS SP8 confocal laser scanning microscope.

### Protein analysis and real time PCR

Protein analysis was performed using Bioplex assays (27-Plex, CXCL-1-, CXCL-2-, CXCL-5-, CCL22-single plex) (Bio-Rad Laboratories) according to the manufacturer’s protocol.

For real time PCR, mRNA was isolated using the InviTrap Spin Universal RNA Mini Kit (Stratec) according to manufacturer’s protocol. mRNA was transcribed into cDNA with Applied Biosytems High Capacity cDNA Reverse Transcription Kit (Thermo Fisher Scientific) according to manufacturer’s protocol. Using Fast Start Universal SYBRGreen Master Rox (Roche) gene expression was measured on Applied Biosystems ViiA7 Real-Time PCR system (Thermo Fisher Scientific) according to manufacturer’s protocol. Primers were ordered from Metabion (www.metabion.com) Sequences can be found in Table S3.

### Statistical analysis

Data of epidermal thickness, Ki67 immunohistochemistry, Bioplex, migration and real time PCR assays were visualized using GraphPad Prism 6 software (https://www.graphpad.com). The unpaired t-test with Welch’s correction was used to test for difference in the means to noRNP sample. The anova test was used to test for differences within the groups for the Ki67 immunohistochemistry assay. Significance level was defined as p < 0.05 (*), p < 0.01 (**), p < 0.001 (***) and p < 0.0001 (****).

## Acknowledgements

We thank Helen Pickersgill from Life Science Editors for editing the manuscript and Jana Sänger as well as Kerstin Weber for excellent technical support. In addition, we thank the NGS core facility at the Helmholtz Center Munich for help with RNA sequencing and the Biobank Biederstein for providing patient samples. RB would like to thank her colleagues at Institute of Computational Biology, Helmholtz Center Munich for helpful discussions.

## Author contributions

Conceptualization: R.B., N.G.S, F.L., N.S.M, S.E., K.E; Methodology: R.B., G.K., M.M., N.S.M, M.J.; Validation M.J.; Formal Analysis R.B., L.C.T., J.E.G.; Investigation: M.J.,; Resources: N.G.S, F.L., C.P., S.R., A.S., A.B., P.S, B. F., J.T.; Data Curation: R.B.; Writing –Original Draft: F.L., N.G.S, S.E., K.E.; Writing – Review & Editing Visualization: R.B, C.B.S-W, T.B., F.T., J.E.G.; Supervision: N.S.M., S.E., K.E.; Project Administration: N.S.M., S.E., K.E.; Funding Acquisition: K.E.

## Supplemental material - figures

**Figure S1:**
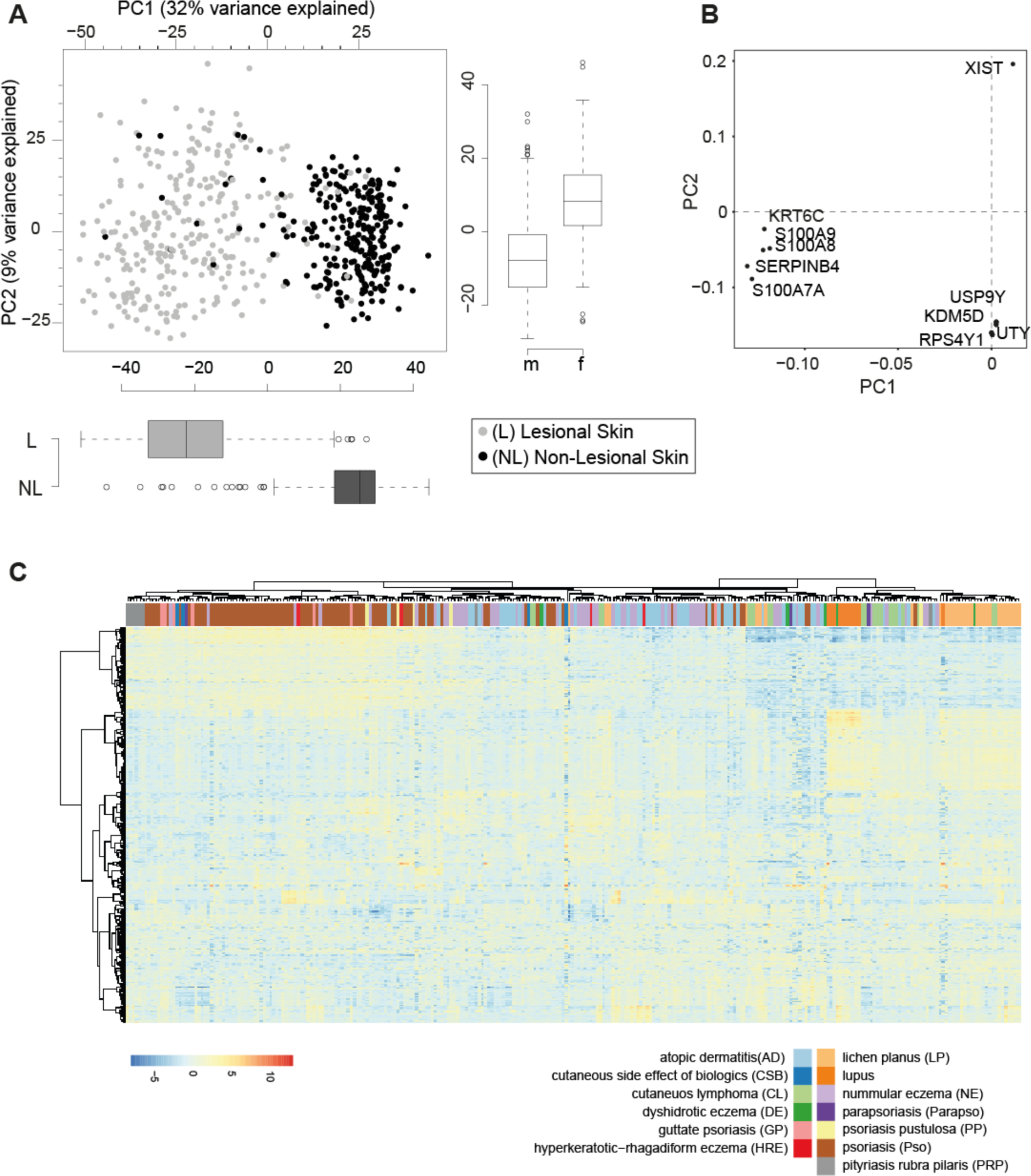
Impact of non-lesional skin and gender on transcriptomics-based clustering. Related to Figure 1. A) Status of the skin (lesional, non-lesional) explains separation of PC1 and in total 32 % of variance in the gene expression data set. The sex (male, female) explains separation of PC2 and 9 % of data set variance. B) Loadings of the principal component analysis depicting the top 10 genes driving PC1 and PC2. C) Hierarchical clustering of gene expression data after correction for gender annotated with diagnosis.

**Figure S2:**
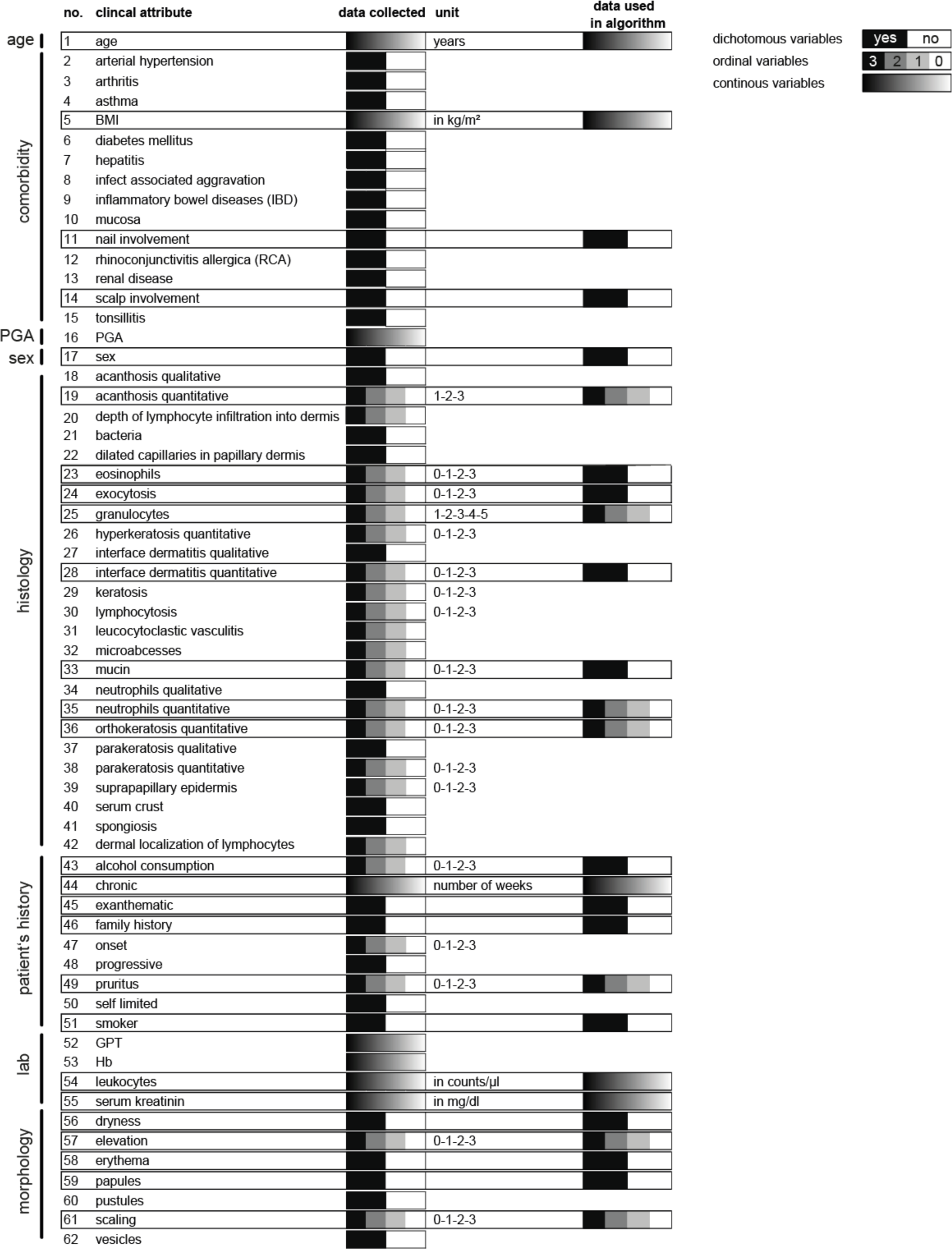
Description of clinical attributes. Related to Figure 1. In total, 62 clinical attributes were obtained from every patient included in the study. These attributes included sex, age, Physician Global Assessment (PGA), information on existing comorbidities (n = 14), histology (n = 23), patient’s history (n = 9), laboratory parameters (lab) (n = 4) and morphology of the skin lesion (n = 7). Data was collected as dichotomous-, ordinal- or continuous variables as indicated. Framed attributes represent the 24 core attributes that were identified and assigned with a gene signature by the AuGER approach.

**Figure S3:**
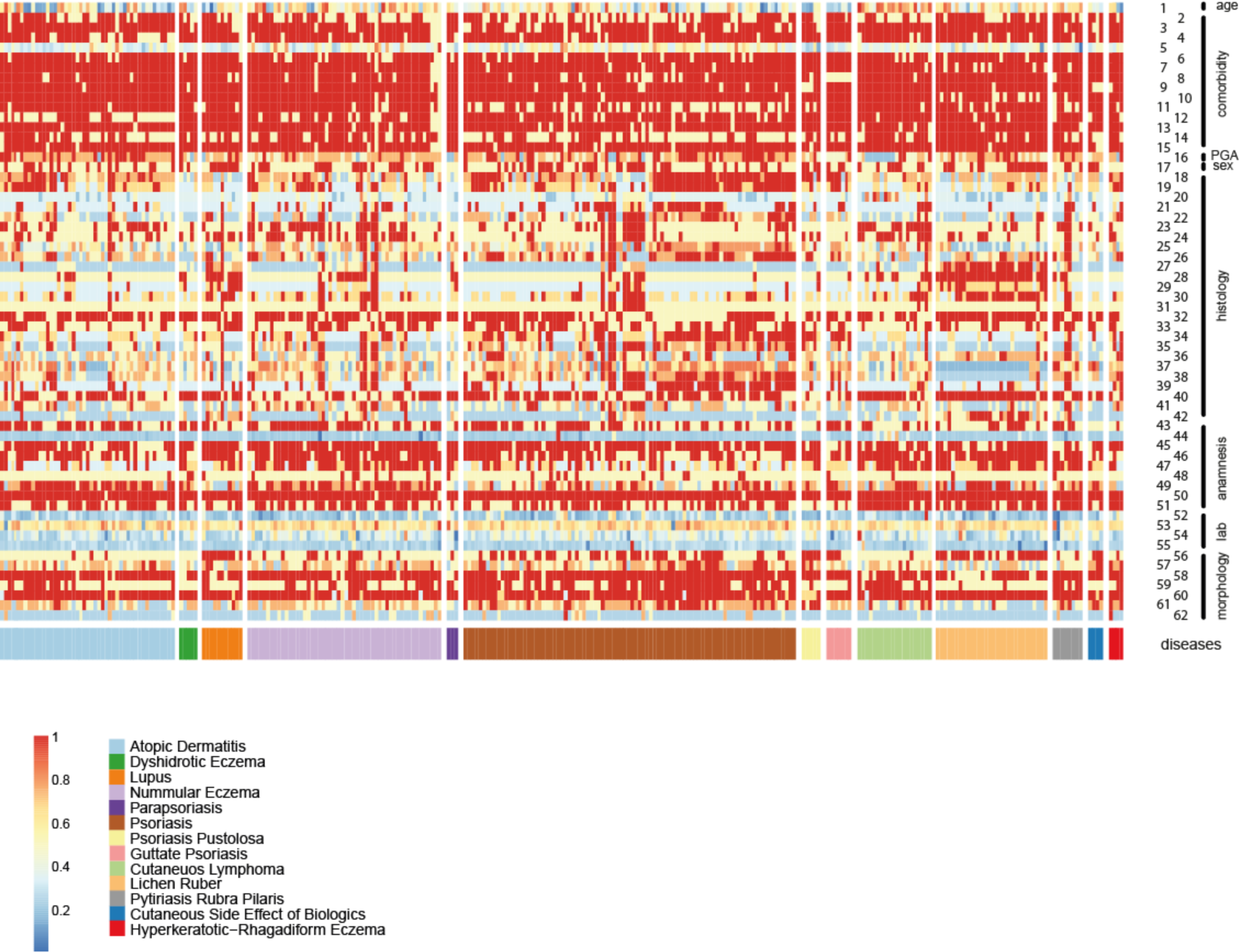
Fingerprints of 287 ncISD patients. Related to Figure 1. Heatmap illustrating each of the 287 patients of the study with his/her personalized clinical phenotype using all 62 attributes as explained in Figure 1B and Figure S2. Patients have been grouped according to their clinical diagnosis as indicated in the legend.

**Figure S4:**
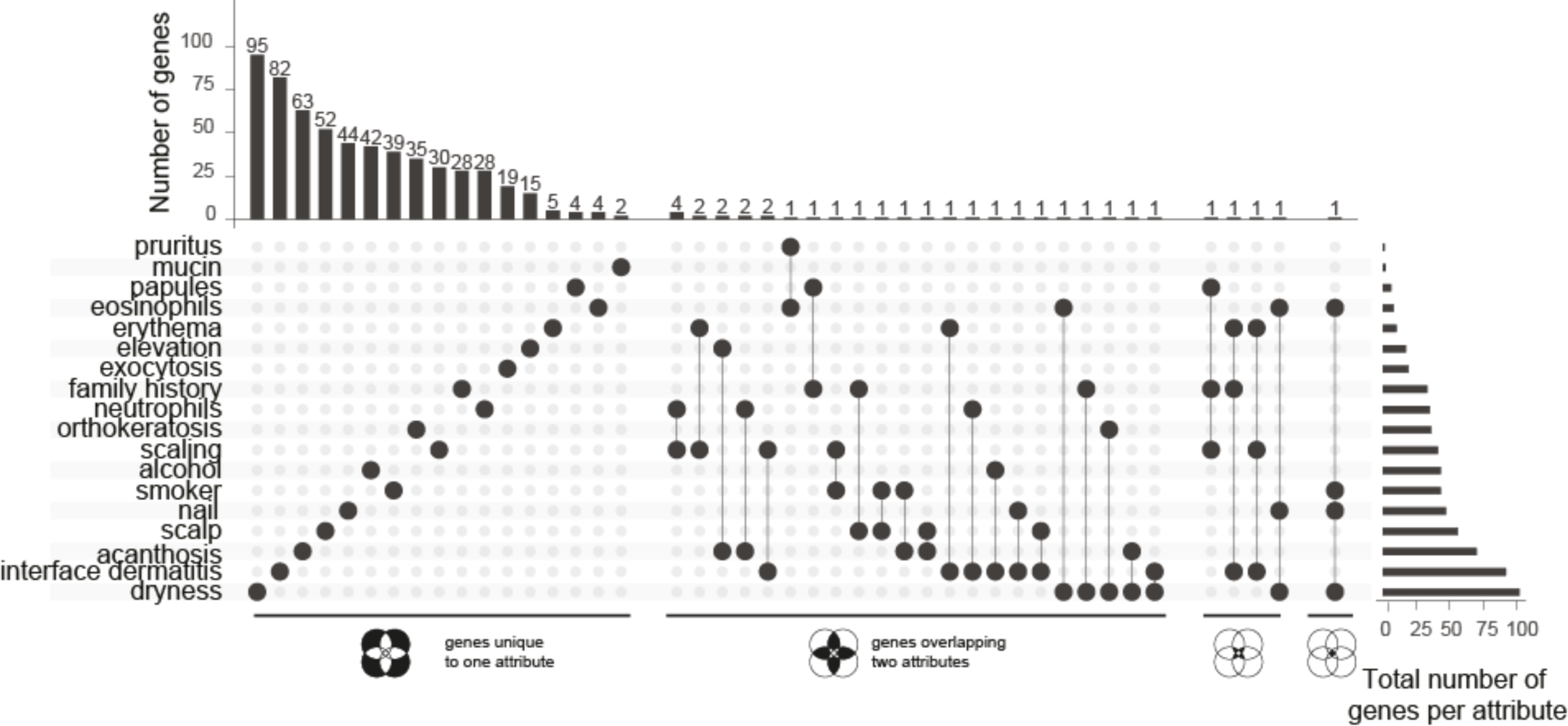
Overlap between the gene sets derived for each attribute using AuGER. Related to Figure 2. The identified core attributes are given on the left and the total number of associated genes by the AuGER approach on the right. On the top, the number of genes are given that are either unique to one attribute (left row), shared between two attributes (middle row), or shared between three or four attributes (right rows). For instance, 102 genes were associated with the attribute ‘dryness’. Of these, 95 genes were unique to ‘dryness’ and the remaining 7 genes were shared with other attributes (with 5 genes shared between ‘dryness’ as well as one other attribute (i.e., ‘eosinophils’, ‘family history’, ‘orthokeratosis’, ‘acanthosis’ or ‘interface dermatitis’); one gene being shared with two other attributes (i.e., ‘eosinophils’ and ‘nail’), and one gene being shared with three other attributes (i.e., ‘eosinophils’, ‘smoker’ and ‘nail’). Overall, 95% of all genes were exclusively associated with one attribute, whereas only 34 genes (5 %) were associated with two, three or at most four attributes simultaneously, corroborating the specificity of the approach.

**Figure S5:**
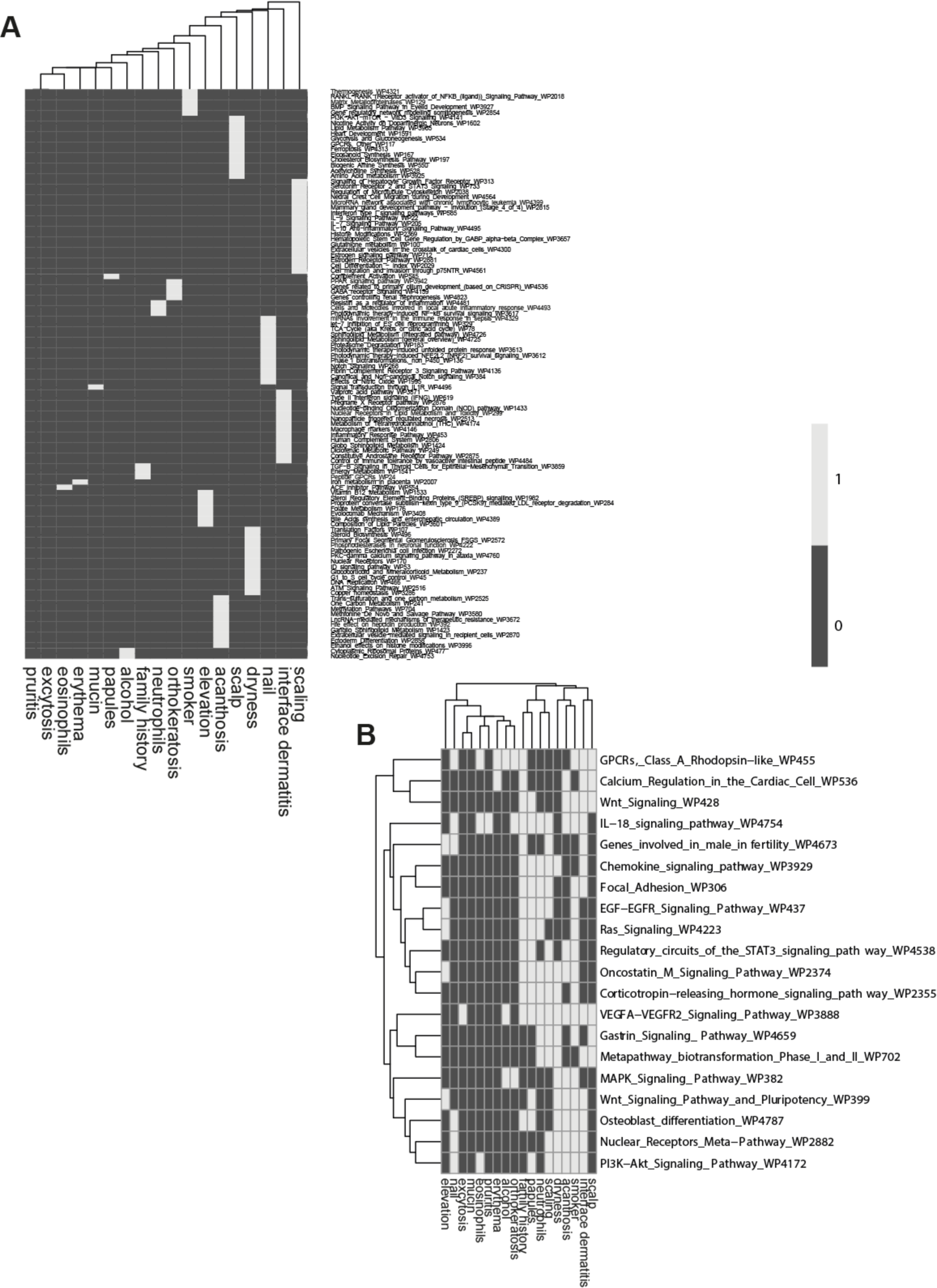
Pathway analysis of each attribute gene signature. Related to Figure 2. A) Unique pathway annotation for each attribute. B) Pathways annotated to five or more attributes. Dark grey (0) indicates no annotation, light grey (1) indicates annotation of an attribute.

**Figure S6:**
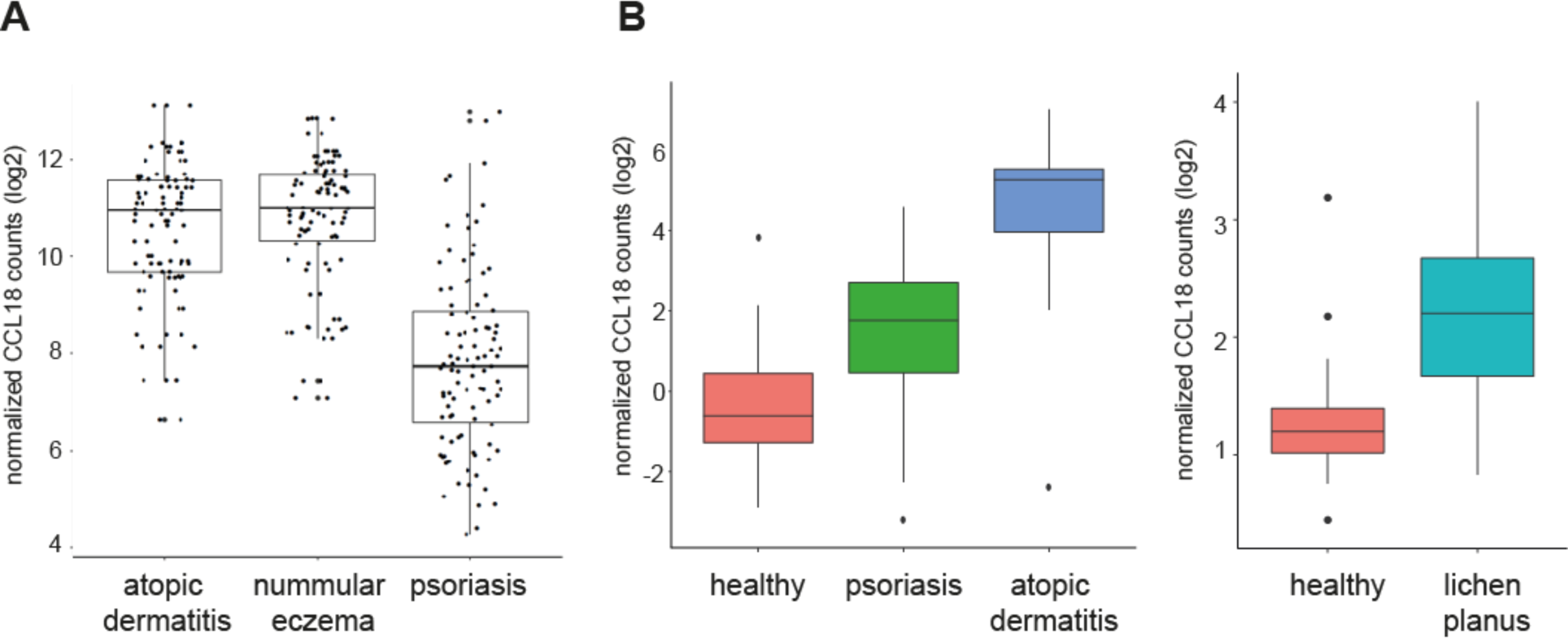
CCL18 levels in lesional skin are associated with pruritic inflammatory skin diseases. Related to Figure 2. A) *CCL18* levels as detected by RNA sequencing from the main cohort of the manuscript, shown for atopic eczema, nummular eczema, and psoriasis patients. B) *CCL18* levels in skin as detected by microarray in a validation cohort of healthy volunteers (n=38) and patients suffering from AE (n=27) and psoriasis (n=37) (left graph) and a RNAseq cohort of healthy volunteers (n=38) and lichen planus (n=20).

**Figure S7:**
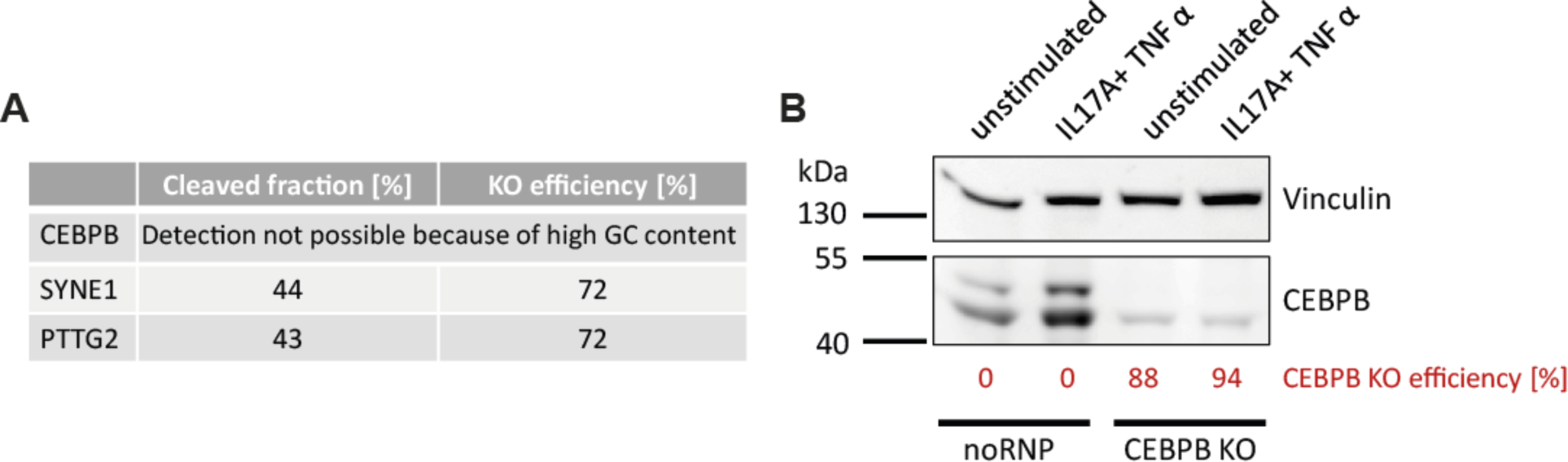
Controls to indicate efficient CRISPR-Cas9 knock-out. Related to figure 4 and 6. A) The SURVEYOR assay was used to determine the knock-out efficiency of SYNE1 and PTTG2 in human primary keratinocytes that have been used for 3D skin models. Due to the high GC content of the CEBPB mRNA, a PCR product could not have been established and analysis by the SURVEYOR assay was not possible. B) Knock-out efficiency of CEBPB determined by western blot analysis of unstimulated and IL-17/TNFa stimulated keratinocytes

**Figure S8:**
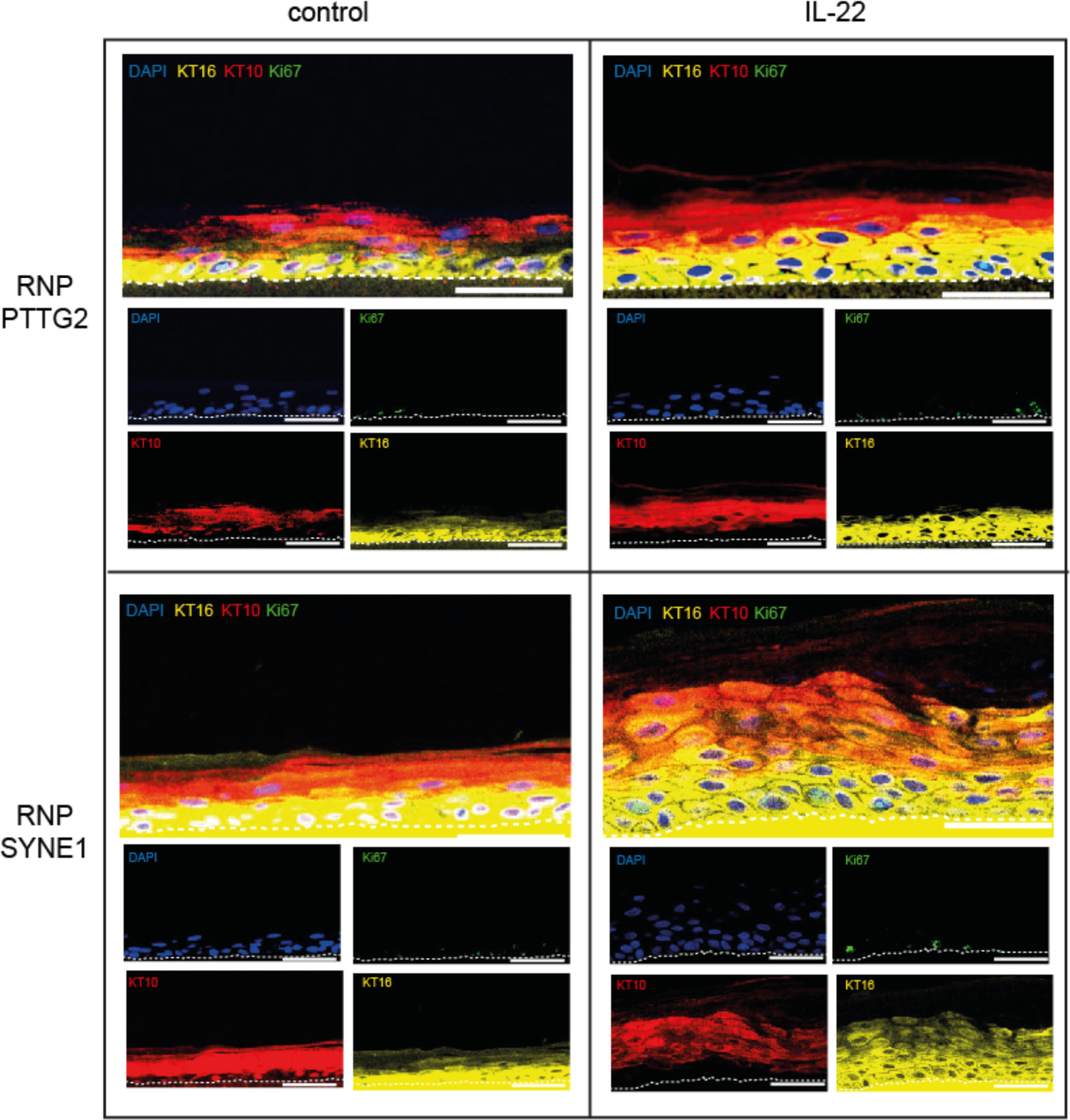
Immunofluorescence staining of 3D skin models. Related to Figure 6. Immunofluorescence staining with Keratin 10 (KT10) (red), Keratin 16 (KT16) (yellow), Ki67 (green) and DAPI (Blue) of 3D skin models with *SYNE1* or *PTTG2* CRISPR-Cas9 knock-out. Acanthosis was induced by stimulation with recombinant IL-22. Unstimulated models served as control. Bars represent 50 µm. Representative Keratin 10 (KT10) (red), Keratin 16 (KT16) (yellow), and Ki67 (green) immunofluorescence staining of 3D skin models with a RNP-CRISPR knock out of *SYNE1* or *PTTG2* with or without (control) IL-22 stimulation. Nuclei were stained with DAPI (blue). The dotted white line indicates the border between basal keratinocytes and the culture membrane. Results on CEBPB are given in Fig. 6 of the main manuscript. Bars represent 50 µm. KT: keratin

## Supplemental material - tables

**Table S1:** Patient characteristica.

| ID | gender | age | diagnosis | BMI | PGA | lesional biopsy | non-lesional biopsy |
| --- | --- | --- | --- | --- | --- | --- | --- |
| 10178 | M | 41,96 | cutaneous lupus | 23 | 3 | x | x |
| 10201 | F | 17,35 | cutaneous lupus | 21,7 | 2 | x | x |
| 10213 | M | 66,03 | cutaneous lupus | 28,1 | 3 | x | x |
| 10217 | F | 68,15 | cutaneous lupus | 25,6 | 3 | x | x |
| 10322 | F | 52,73 | cutaneous lupus | 18,2 | 2 | x |  |
| 10359 | F | 69,03 | cutaneous lupus | 18,6 | 3 | x | x |
| 10381 | F | 36,01 | cutaneous lupus | 22,8 | 3 | x | x |
| 10409 | F | 59,80 | cutaneous lupus | 20,8 | 4 | x | x |
| 10418 | M | 26,94 | cutaneous lupus | 27,8 | 1 | x | x |
| 10421 | F | 44,47 | cutaneous lupus | 23,3 | 4 | x | x |
| 10710 | F | 54,30 | cutaneous lupus | 24,2 | 3 | x | x |
| *n=11* | *F: 8; M: 3* | *48.8*<br>*±17.16* |  | *23.1*<br>*±3.26* | *2.82*<br>*±0.87* |  |  |
| 10005 | M | 44,79 | lichen planus | 27,7 | 3 | x | x |
| 10009 | F | 79,30 | lichen planus | 23,6 | 3 | x | x |
| 10016 | F | 43,92 | lichen planus | 18,4 | 4 | x | x |
| 10053 | M | 51,07 | lichen planus |  | 3 | x | x |
| 10069 | F | 65,30 | lichen planus | 27 | 4 | x | x |
| 10074 | M | 71,55 | lichen planus |  | 2 | x | x |
| 10075 | F | 51,70 | lichen planus |  | 2 | x | x |
| 10076 | F | 75,48 | lichen planus |  | 2 | x | x |
| 10104 | M | 43,22 | lichen planus |  | 2 | x | x |
| 10109 | M | 59,93 | lichen planus | 32 | 4 | x | x |
| 10113 | M | 65,17 | lichen planus | 24,6 | 3 | x | x |
| 10174 | F | 39,04 | lichen planus | 19,4 | 3 | x | x |
| 10212 | F | 74,00 | lichen planus | 24 | 2 | x | x |
| 10317 | M | 61,83 | lichen planus | 32,6 | 3 | x | x |
| 10318 | F | 54,53 | lichen planus | 25,2 | 3 | x | x |
| 10340 | F | 65,22 | lichen planus | 28,2 | 4 | x | x |
| 10347 | F | 78,65 | lichen planus | 25,7 | 4 | x | x |
| 10382 | M | 63,89 | lichen planus | 36,8 | 2 | x | x |
| 10422 | M | 54,78 | lichen planus | 34 | 3 | x | x |
| 10435 | F | 33,83 | lichen planus | 21,1 | 4 | x | x |
| 10458 | F | 57,63 | lichen planus |  | 3 | x | x |
| 10492 | F | 84,65 | lichen planus | 22,9 | 2 | x | x |
| 10496 | M | 56,15 | lichen planus | 30 | 4 | x | x |
| 10532 | F | 70,61 | lichen planus | 28,7 | 2 | x | x |
| 10576 | F | 54,70 | lichen planus | 32,8 | 3 | x | x |
| 10613 | M | 28,92 | lichen planus | 24,2 | 3 | x | x |
| 10622 | M | 72,18 | lichen planus | 25,5 | 3 | x | x |
| 10640 | F | 54,00 | lichen planus | 26,2 | 3 | x |  |
| 10687 | F | 65,83 | lichen planus | 25,4 | 2 | x | x |
| 10705 | M | 43,18 | lichen planus | 22,2 | 3 | x | x |
| *n=30* | *F: 17; M: 13* | *58.83*<br>*±14* |  | *26.59*<br>*±4.64* | *2.93*<br>*±0.74* |  |  |
| 10004 | F | 27,85 | AD | 19,4 | 4 | x | x |
| 10007 | M | 27,48 | AD | 19 | 4 | x | x |
| 10020 | F | 22,49 | AD | 20 | 3 | x | x |
| 10025 | M | 23,61 | AD | 22,7 | 4 | x | x |
| 10043 | M | 72,33 | AD | 19,8 | 4 | x | x |
| 10063 | F | 27,10 | AD | 21,97 | 3 | x | x |
| 10085 | F | 35,09 | AD | 32 | 4 | x | x |
| 10089 | M | 62,30 | AD |  | 3 | x | x |
| 10137 | M | 29,01 | AD | 25,8 | 4 | x | x |
| 10146 | M | 25,89 | AD | 21 | 3 | x | x |
| 10248 | F | 53,13 | AD | 27,8 |  | x | x |
| 10264 | M | 57,30 | AD |  | 3 | x | x |
| 10269 | M | 54,18 | AD | 21 |  | x | x |
| 10294 | M | 50,01 | AD | 24,6 | 3 | x | x |
| 10321 | M | 79,55 | AD |  | 3 | x | x |
| 10393 | M | 39,44 | AD | 32,6 | 4 | x | x |
| 10401 | M | 77,67 | AD | 20,8 | 3 | x | x |
| 10416 | M | 61,81 | AD | 30 | 4 | x | x |
| 10442 | M | 22,75 | AD | 34,1 | 3 | x | x |
| 10452 | M | 51,21 | AD | 20,1 | 3 | x | x |
| 10462 | F | 66,44 | AD | 24,7 | 3 | x | x |
| 10655 | M | 53,69 | AD | 32,1 |  | x | x |
| 10671 | F | 23,90 | AD | 22,1 |  | x |  |
| 10682 | F | 21,50 | AD | 26,6 |  | x | x |
| 13237 | M | 56,01 | AD | 30,1 | 4 | x | x |
| 16083 | M | 31,45 | AD | 24,57 | 4 | x | x |
| 24627 | F | 47,59 | AD | 34,19 | 3 | x | x |
| 28093 | F | 26,15 | AD | 21,33 | 3 | x | x |
| 37685 | M | 38,74 | AD | 20,2 | 3 | x | x |
| 48724 | M | 49,59 | AD | 21,96 | 4 | x | x |
| 51694 | M | 55,98 | AD | 30,75 | 3 | x | x |
| 57856 | M | 36,77 | AD | 23,95 | 3 | x | x |
| 71982 | M | 41,64 | AD | 29,72 | 3 | x | x |
| 83239 | M | 33,07 | AD | 15,56 | 4 | x | x |
| 10464 | M | 56,69 | AD | 23,5 | 3 | x | x |
| 10570 | F | 45,04 | AD | 31,7 |  | x | x |
| 10367 | M | 68,47 | AD | 21,2 | 4 | x | x |
| 10035 | F | 51,51 | AD | 45,7 | 4 | x | x |
| 10071 | F | 79,26 | AD | 23,5 | 4 | x | x |
| 10091 | M | 81,55 | AD |  | 2 | x | x |
| 10147 | M | 25,31 | AD | 23 | 3 | x | x |
| 10149 | F | 87,00 | AD | 23 | 4 | x | x |
| 10175 | M | 23,22 | AD | 27 | 3 | x | x |
| 10320 | M | 47,50 | AD | 30,1 | 3 | x | x |
| 10498 | M | 74,21 | AD | 26,5 | 3 | x | x |
| 10654 | M | 56,75 | AD | 28,7 |  | x | x |
| 10664 | F | 73,99 | AD | 23 |  | x | x |
| *n=47* | *F: 15; M: 32* | *47.94*<br>*±19.22* |  | *25.52*<br>*±5.63* | *3.38*<br>*±0.54* |  |  |
| 10166 | F | 79,27 | DE | 26 | 4 | x | x |
| 10223 | M | 82,09 | DE |  | 3 | x | x |
| 10410 | M | 52,49 | DE | 27,8 | 3 | x | x |
| *n=3* | *F:1, M: 2* | *71.28*<br>*±16.33* |  | *26.9*<br>*±1.27* | *2.22*<br>*±0.58* |  |  |
| 10028 | M | 66,52 | HRE | 21,6 | 3 | x | x |
| 10073 | M | 21,93 | HRE | 28,4 | 4 | x | x |
| 10082 | M | 29,72 | HRE | 32,7 | 1 | x | x |
| *n=3* | *F:0, M: 3* | *39.39*<br>*±23.81* |  | *27.57*<br>*±5.6* | *2.67*<br>*±1.53* |  |  |
| 10021 | F | 66,20 | nummular eczema |  | 4 | x | x |
| 10084 | F | 51,72 | nummular eczema |  | 2 | x | x |
| 10144 | F | 55,46 | nummular eczema | 22,3 | 3 | x | x |
| 10150 | M | 67,60 | nummular eczema | 23 | 2 | x | x |
| 10165 | M | 60,35 | nummular eczema | 18 | 4 | x | x |
| 10220 | M | 35,70 | nummular eczema | 44,2 | 4 | x | x |
| 10309 | M | 78,82 | nummular eczema | 31,5 | 2 | x | x |
| 10314 | F | 71,42 | nummular eczema |  | 2 | x | x |
| 10355 | F | 87,65 | nummular eczema | 29,7 | 3 | x | x |
| 10361 | F | 38,99 | nummular eczema | 29,8 | 3 | x | x |
| 10387 | M | 66,60 | nummular eczema | 45,6 | 2 | x | x |
| 10389 | M | 49,12 | nummular eczema | 19,6 | 3 | x | x |
| 10400 | M | 71,40 | nummular eczema | 27 | 3 | x | x |
| 10404 | F | 67,17 | nummular eczema | 25,9 | 3 | x | x |
| 10406 | M | 58,18 | nummular eczema | 30,7 | 3 | x | x |
| 10413 | M | 33,40 | nummular eczema | 32 | 3 | x | x |
| 10417 | M | 74,37 | nummular eczema | 29,4 | 3 | x | x |
| 10426 | F | 81,35 | nummular eczema | 27,1 | 3 | x | x |
| 10443 | M | 79,37 | nummular eczema | 27,6 | 3 | x | x |
| 10446 | F | 79,82 | nummular eczema | 23,2 | 3 | x | x |
| 10450 | F | 69,04 | nummular eczema | 27 | 3 | x | x |
| 10453 | F | 89,89 | nummular eczema | 20,5 | 4 | x | x |
| 10455 | M | 63,95 | nummular eczema | 42,6 | 3 | x | x |
| 10466 | F | 29,63 | nummular eczema | 18,4 | 2 | x | x |
| 10479 | F | 59,07 | nummular eczema | 31,1 | 3 | x | x |
| 10481 | M | 24,57 | nummular eczema | 20,7 | 3 | x | x |
| 10483 | F | 77,36 | nummular eczema | 23,1 | 2 | x | x |
| 10528 | M | 54,09 | nummular eczema | 34 | 3 | x | x |
| 10535 | M | 68,29 | nummular eczema | 23,8 | 3 | x | x |
| 10539 | M | 63,14 | nummular eczema | 29,4 | 2 | x | x |
| 10544 | F | 48,98 | nummular eczema |  | 2 | x | x |
| 10558 | M | 90,76 | nummular eczema | 22,1 |  | x | x |
| 10573 | F | 40,55 | nummular eczema | 29,7 | 2 | x | x |
| 10577 | F | 83,80 | nummular eczema | 22,6 | 3 | x | x |
| 10580 | M | 28,11 | nummular eczema | 21,7 | 3 | x | x |
| 10581 | M | 60,12 | nummular eczema | 27,8 | 3 | x | x |
| 10592 | M | 35,25 | nummular eczema | 26,5 | 3 | x | x |
| 10596 | F | 54,38 | nummular eczema |  | 3 | x | x |
| 10598 | M | 52,50 | nummular eczema |  | 3 | x | x |
| 10599 | M | 69,40 | nummular eczema |  | 3 | x | x |
| 10601 | M | 31,25 | nummular eczema | 40,23 |  | x | x |
| 10610 | F | 67,99 | nummular eczema | 31,6 |  | x | x |
| 10624 | M | 82,92 | nummular eczema | 25,3 | 3 | x | x |
| 10634 | F | 64,78 | nummular eczema |  | 3 | x | x |
| 10636 | M | 72,95 | nummular eczema |  | 3 | x | x |
| 10637 | M | 63,49 | nummular eczema | 32,3 | 3 | x | x |
| 10643 | M | 55,30 | nummular eczema |  |  | x | x |
| 10644 | F | 70,49 | nummular eczema | 25,2 |  | x | x |
| 10659 | F | 71,22 | nummular eczema | 26,9 |  | x | x |
| 10670 | M | 70,32 | nummular eczema | 25 |  | x | x |
| 10685 | M | 54,87 | nummular eczema |  | 3 | x | x |
| *n=51* | *F: 22,*<br>*M:29* | *61.63*<br>*±16.87* |  | *27.85*<br>*±6.59* | *2.86*<br>*±0.55* |  |  |
| 10366 | M | 82,30 | cut. lymphoma | 24,2 | 3 | x | x |
| 10390 | F | 85,65 | cut. lymphoma |  | 4 | x | x |
| 10392 | F | 92,91 | cut. lymphoma | 31,2 | 2 | x | x |
| 10402 | M | 84,29 | cut. lymphoma | 25,5 | 1 | x | x |
| 10407 | M | 75,51 | cut. lymphoma |  | 4 | x | x |
| 10430 | M | 52,15 | cut. lymphoma | 27,7 | 1 | x | x |
| 10440 | M | 70,38 | cut. lymphoma | 25,8 | 2 | x | x |
| 10467 | M | 61,97 | cut. lymphoma |  | 2 | x | x |
| 10504 | M | 88,62 | cut. lymphoma | 29,9 | 2 | x | x |
| 10549 | F | 77,23 | cut. lymphoma | 25,3 | 3 | x | x |
| HG16.1069 | F | 86,53 | cut. lymphoma | 24,2 | 1 | x | x |
| HG16.1099 | M | 73,49 | cut. lymphoma | 22,7 | 2 | x | x |
| HG16.1628 | M | 46,38 | cut. lymphoma | 24 | 1 | x | x |
| HG16.537 | M | 68,15 | cut. lymphoma | 22 | 1 | x | x |
| HG16.668 | M | 63,35 | cut. lymphoma | 22,1 | 1 | x | x |
| HG16.711 | F | 70,83 | cut. lymphoma | 23,4 | 2 | x | x |
| HG16.734 | F | 82,26 | cut. lymphoma | 29 | 1 | x | x |
| HG16.975 | F | 78,57 | cut. lymphoma | 23 | 1 | x | x |
| 10607 | F | 43,13 | cut. lymphoma | 25,4 | 3 | x | x |
| 10658 | M | 86,45 | cut. lymphoma | 23,7 | 4 | x |  |
| *n=20* | *F: 8; M:*<br>*12* | *73.51*<br>*±14.14* |  | *25.24*<br>*±2.73* | *2.05*<br>*±1.09* |  |  |
| 10283 | M | 36,69 | CSB | 23,9 | 1 | x | x |
| 10371 | F | 39,96 | CSB | 21,9 | 2 | x | x |
| 10349 | M | 65,52 | CSB | 24,4 | 3 | x | x |
| 10632 | M | 45,19 | CSB | 29 |  | x | x |
| 10353 | F | 44,66 | CSB |  | 3 | x | x |
| 10344 | F | 58,66 | CSB |  | 2 | x | x |
| 10346 | F | 30,58 | CSB |  | 3 | x | x |
| 10383 | M | 25,42 | CSB | 19 | 3 | x | x |
| 10388 | F | 28,05 | CSB | 17 | 2 | x | x |
| 10420 | M | 41,25 | CSB | 29,3 | 3 | x | x |
| 10447 | F | 57,53 | CSB |  | 2 | x | x |
| 10541 | F | 47,44 | CSB | 30 | 3 | x | x |
| *n=12* | *F: 7; M: 5* | *43.41*<br>*±12.53* |  | *24.31*<br>*±4.88* | *2.45*<br>*±0.69* |  |  |
| 10002 | F | 18,33 | psoriasis | 26,8 | 4 | x | x |
| 10006 | M | 42,72 | psoriasis | 37 | 4 | x | x |
| 10013 | M | 50,26 | psoriasis | 33,6 | 4 | x | x |
| 10030 | M | 38,26 | psoriasis | 30,6 | 4 | x | x |
| 10033 | M | 67,58 | psoriasis | 32,9 | 4 | x | x |
| 10034 | F | 67,48 | psoriasis | 29,6 | 4 | x | x |
| 10036 | M | 52,63 | psoriasis | 36,7 | 4 | x | x |
| 10046 | M | 54,78 | psoriasis | 32,2 | 3 | x | x |
| 10049 | M | 45,79 | psoriasis | 22,8 | 3 | x | x |
| 10051 | F | 42,20 | psoriasis | 23,2 | 3 | x | x |
| 10052 | F | 90,62 | psoriasis | 33,2 | 3 | x | x |
| 10056 | M | 22,99 | psoriasis | 22,6 | 3 | x | x |
| 10058 | F | 70,31 | psoriasis | 43,7 | 4 | x | x |
| 10060 | M | 52,93 | psoriasis | 24 | 4 | x | x |
| 10067 | M | 42,30 | psoriasis | 21,9 | 3 | x | x |
| 10072 | F | 31,69 | psoriasis | 22 | 3 | x | x |
| 10078 | M | 51,36 | psoriasis | 26,1 |  | x |  |
| 10079 | M | 24,72 | psoriasis |  | 3 | x | x |
| 10083 | F | 41,79 | psoriasis | 40,3 | 4 | x | x |
| 10103 | M | 55,23 | psoriasis | 31,7 | 4 | x | x |
| 10119 | F | 57,01 | psoriasis | 23 | 4 | x | x |
| 10128 | F | 53,49 | psoriasis | 30 | 3 | x | x |
| 10133 | M | 63,70 | psoriasis | 31,2 | 3 | x | x |
| 10138 | F | 68,62 | psoriasis | 21 | 3 | x | x |
| 10153 | F | 46,87 | psoriasis | 33 | 3 | x | x |
| 10154 | F | 55,43 | psoriasis | 47 | 4 | x | x |
| 10158 | M | 55,42 | psoriasis | 24,6 | 2 | x | x |
| 10171 | M | 56,98 | psoriasis | 31 | 2 | x | x |
| 10172 | F | 90,97 | psoriasis | 26 | 2 | x | x |
| 10195 | M | 47,40 | psoriasis | 21,4 | 3 | x | x |
| 10198 | M | 54,32 | psoriasis | 29 | 3 | x | x |
| 10199 | F | 44,12 | psoriasis | 34 | 4 | x | x |
| 10200 | M | 38,52 | psoriasis | 32 | 4 | x | x |
| 10203 | M | 49,74 | psoriasis | 26 | 4 | x | x |
| 10206 | F | 27,58 | psoriasis | 29 | 3 | x | x |
| 10208 | F | 65,75 | psoriasis | 26,7 | 3 | x | x |
| 10210 | F | 50,70 | psoriasis |  | 2 | x | x |
| 10227 | M | 54,64 | psoriasis | 29,3 | 3 | x | x |
| 10233 | M | 58,41 | psoriasis | 23,1 | 4 | x | x |
| 10239 | F | 88,72 | psoriasis | 21,9 | 4 | x | x |
| 10242 | F | 65,87 | psoriasis | 34,7 |  | x | x |
| 10249 | M | 72,65 | psoriasis | 28,7 |  | x | x |
| 10282 | M | 38,66 | psoriasis | 26,3 | 1 | x | x |
| 10297 | F | 66,59 | psoriasis | 26,4 | 2 | x | x |
| 10298 | M | 71,42 | psoriasis | 29,6 | 3 | x | x |
| 10307 | M | 46,40 | psoriasis | 32,2 | 4 | x | x |
| 10327 | M | 44,07 | psoriasis | 32,7 | 2 | x | x |
| 10343 | M | 76,58 | psoriasis |  | 3 | x | x |
| 10354 | M | 74,73 | psoriasis | 23,7 | 4 | x | x |
| 10384 | M | 47,07 | psoriasis | 39,1 | 4 | x | x |
| 10429 | M | 60,89 | psoriasis |  | 3 | x | x |
| 10456 | M | 33,87 | psoriasis | 29 | 3 | x | x |
| 10465 | M | 49,82 | psoriasis | 23,8 | 3 | x | x |
| 10489 | F | 84,55 | psoriasis |  | 2 | x | x |
| 10542 | M | 57,34 | psoriasis |  |  | x |  |
| 10562 | F | 42,45 | psoriasis | 23,2 |  | x | x |
| 10574 | M | 39,34 | psoriasis | 20,8 |  | x | x |
| 10578 | M | 51,88 | psoriasis | 29,4 |  | x | x |
| 10590 | M | 61,66 | psoriasis | 29,9 |  | x | x |
| 10602 | M | 55,67 | psoriasis | 21,6 |  | x | x |
| 10609 | F | 73,35 | psoriasis | 29,8 |  | x | x |
| 10615 | M | 66,90 | psoriasis | 26,7 |  | x | x |
| 10620 | M | 36,87 | psoriasis | 29,8 |  | x | x |
| 10630 | F | 58,20 | psoriasis | 21,5 |  | x | x |
| 10646 | M | 58,55 | psoriasis | 26 |  | x | x |
| 10649 | M | 53,56 | psoriasis | 25,4 |  | x | x |
| 10650 | F | 43,52 | psoriasis | 29,7 |  | x | x |
| 10656 | F | 48,49 | psoriasis | 27,1 |  | x | x |
| 10665 | M | 25,84 | psoriasis | 32,4 |  | x | x |
| 10668 | F | 25,09 | psoriasis | 26,4 |  | x | x |
| 10691 | M | 46,75 | psoriasis |  |  | x | x |
| 10350 | F | 73,52 | psoriasis |  | 3 | x | x |
| 10419 | F | 73,67 | psoriasis |  | 3 | x | x |
| 10064 | F | 92,21 | psoriasis | 31,6 | 2 | x | x |
| 10136 | F | 67,50 | psoriasis | 30 | 3 | x | x |
| 10626 | M | 68,13 | psoriasis | 22 |  | x | x |
| 10177 | M | 51,57 | psoriasis | 34 | 2 | x | x |
| 10369 | F | 51,61 | psoriasis | 17,1 | 3 | x | x |
| 10372 | M | 78,22 | psoriasis | 35,5 | 3 | x | x |
| 10379 | F | 68,84 | psoriasis | 36,3 | 3 | x | x |
| 10482 | F | 58,82 | psoriasis |  | 3 | x | x |
| 10631 | M | 24,64 | psoriasis | 35,6 |  | x | x |
| 10633 | F | 37,32 | psoriasis | 24,8 |  | x | x |
| 10173 | F | 56,78 | psoriasis | 36 | 4 | x | x |
| 10211 | M | 39,41 | psoriasis | 40,35 | 4 | x | x |
| 10616 | M | 67,34 | psoriasis | 27,8 | 4 | x | x |
| 10645 | F | 59,98 | psoriasis | 32,9 |  | x | x |
| 10019 | M | 59,32 | psoriasis | 31,1 | 4 | x | x |
| 10159 | F | 53,41 | psoriasis | 36,3 | 4 | x | x |
| 10386 | M | 81,76 | psoriasis | 22,3 | 2 | x | x |
| 10461 | F | 53,80 | psoriasis | 48 | 3 | x | x |
| 10235 | F | 42,82 | psoriasis | 20,9 | 1 | x | x |
| *n= 92* | *F: 40; M: 52* | 54.69<br>*±16.05* |  | 29.25<br>*±6.14* | 3.18<br>*±0.79* |  |  |
| 10255 | M | 46,76 | Parapsoriasis |  | 3 | x | x |
| 10439 | M | 67,28 | Parapsoriasis | 27,2 | 1 | x | x |
| 10449 | M | 57,99 | Parapsoriasis | 27,5 | 1 | x | x |
| *n= 3* | *F: 0; M: 3* | 57.34<br>*±10.27* |  | 27.35<br>*±0.21* | 1.67<br>*±1.15* |  |  |
| 10087 | F | 32,33 | pso guttata | 21 | 2 | x | x |
| 10107 | F | 48,88 | pso guttata | 21,2 | 2 | x | x |
| 10205 | M | 46,44 | pso guttata | 26 | 3 | x | x |
| 10291 | M | 37,92 | pso guttata | 25,5 | 2 | x | x |
| 10296 | M | 34,21 | pso guttata |  | 2 | x | x |
| 10471 | F | 34,98 | pso guttata | 19,3 | 2 | x | x |
| 10276 | M | 28,77 | pso guttata |  | 4 | x | x |
| *n=7* | *F: 3; M: 4* | 37.65<br>*±7.41* |  | 22.6<br>*±2.97* | 2.43<br>*±0.79* |  |  |
| 10086 | M | 65,12 | PRP |  | 3 | x | x |
| 10478 | M | 77,36 | PRP | 26,1 | 4 | x | x |
| 10524 | F | 76,73 | PRP |  | 3 | x | x |
| 10561 | F | 65,12 | PRP | 28 | 3 | x | x |
| 10648 | M | 69,69 | PRP | 19,7 | 2,5 | x |  |
| 10666 | M | 62,00 | PRP | 24,8 | 3 | x | x |
| 10690 | M | 65,26 | PRP | 25,7 | 4 | x | x |
| 10721 | F | 46,38 | PRP | 28,7 | 3 | x | x |
| *n=8* | *F: 2; M:5* | 65.96<br>*±9.72* |  | 25.5<br>*±3.19* | 3.19<br>*±0.53* |  |  |

**Table S2:**
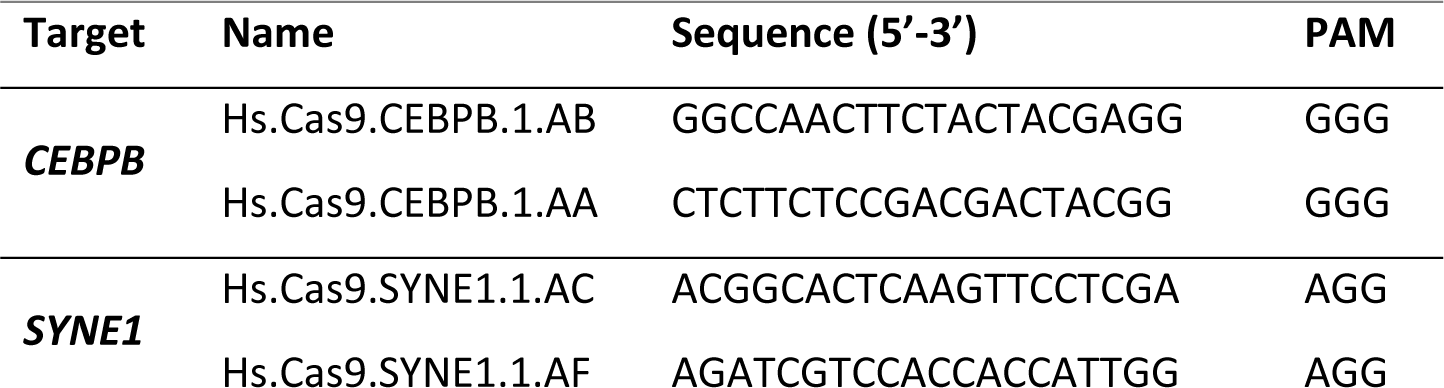
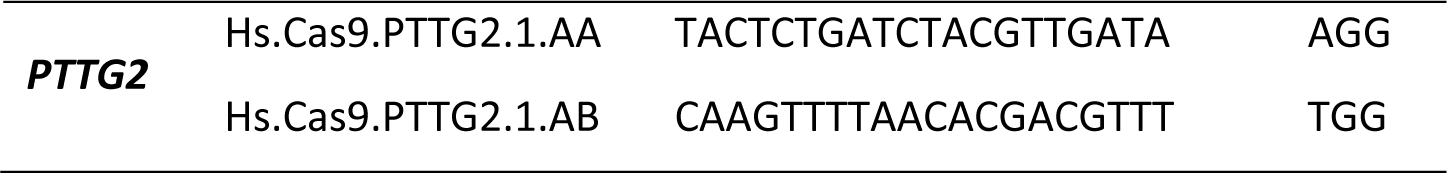
crRNA sequences.

**Table S3:**
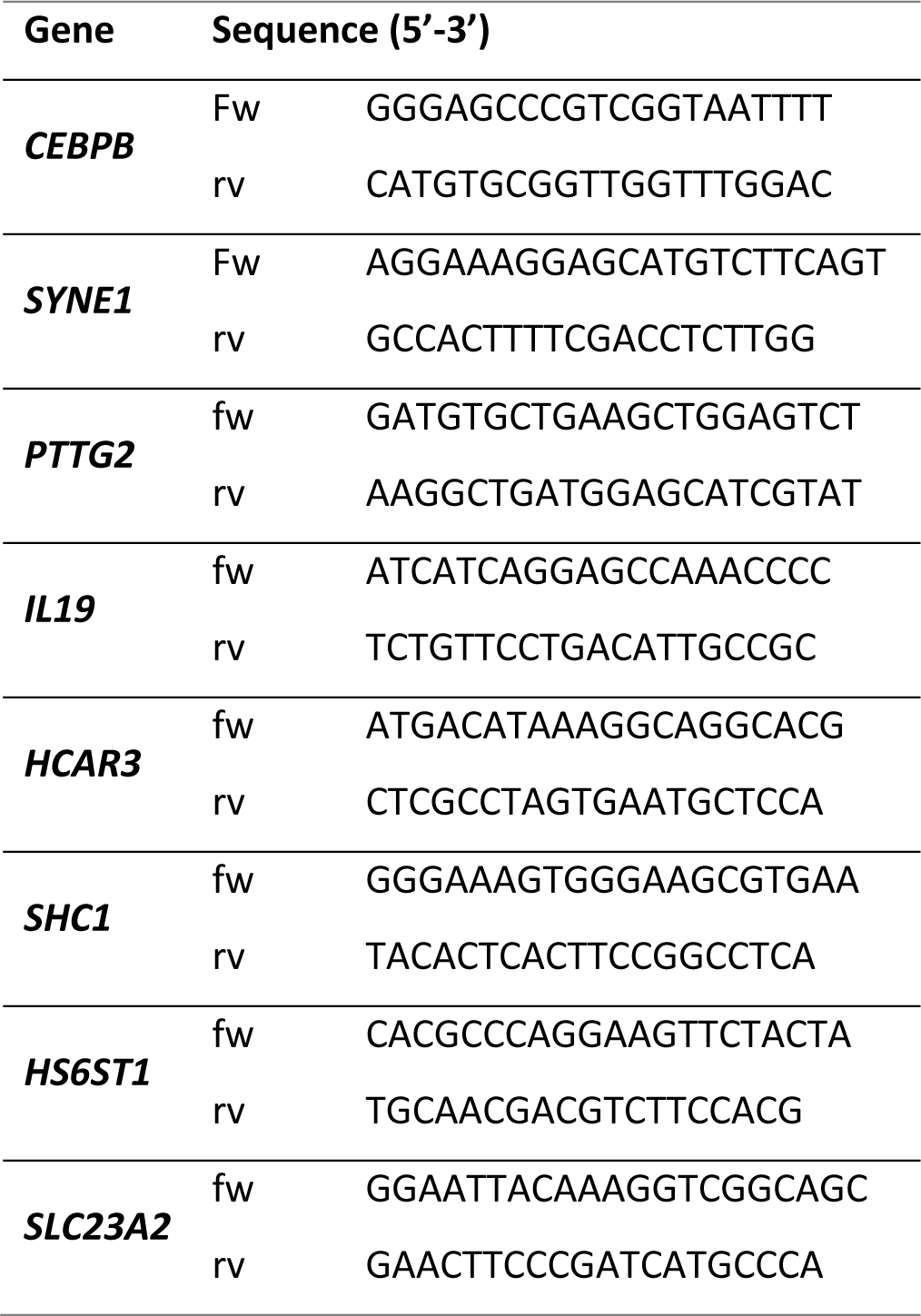
Primer sequences.

**Table S4:**
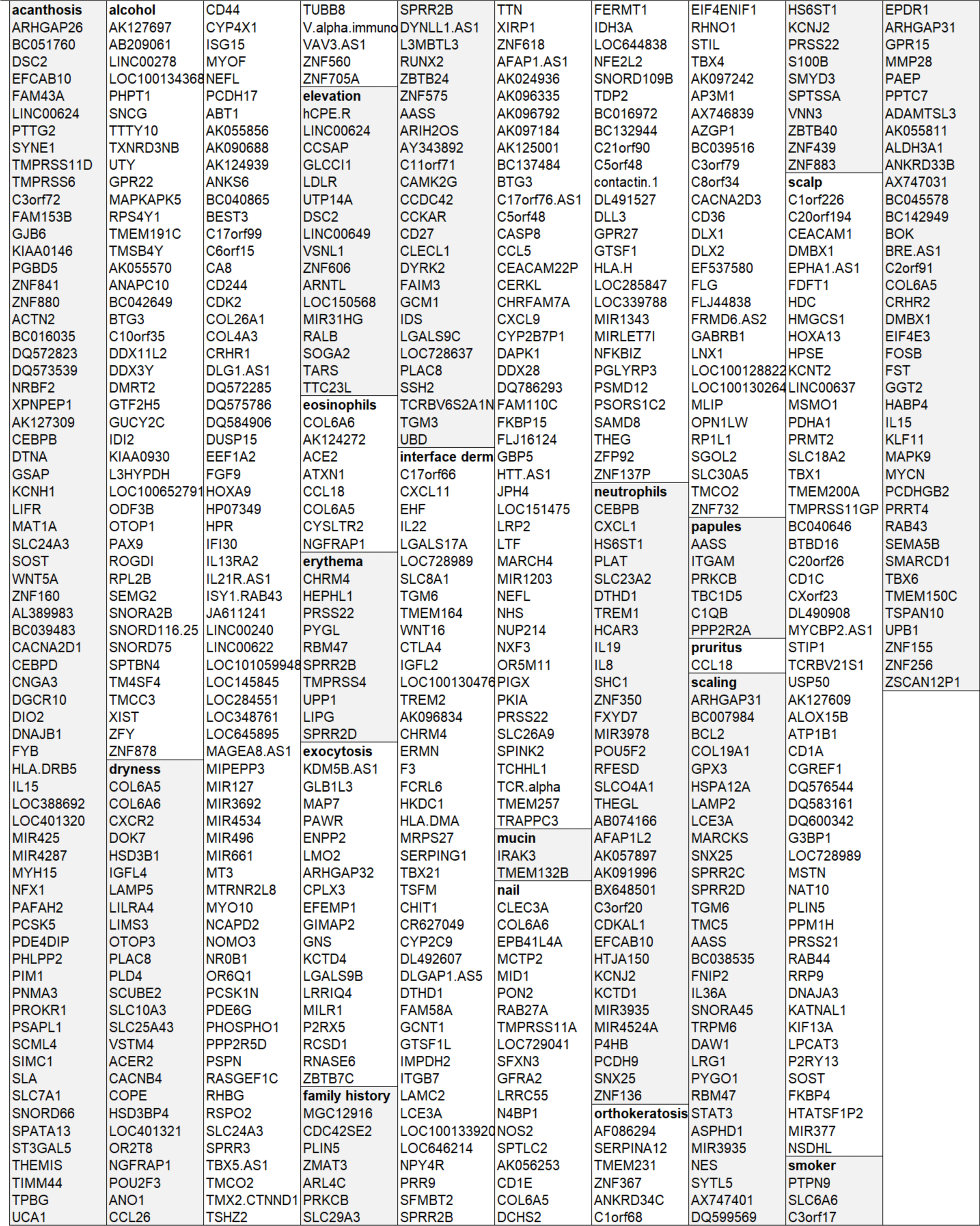
Gene lists of each core attribute-gene association.

